# PROTEOME DATA BASED IDENTIFICATION OF POTENTIAL RNAi TARGETS FOR COTTON MEALYBUG (*Phenacoccus solenopsis* Tinsley) POPULATION MANAGEMENT

**DOI:** 10.1101/2024.03.08.584030

**Authors:** Sanchita Singh, Somnath Rahangdale, Shivali Pandita, Manisha Singh, Gauri Saxena, Gaurav Jain, Praveen C. Verma

## Abstract

**Background *of the study*:** *Phenacoccus solenopsis* Tinsley (Hemiptera: Pseudococcidae), commonly known as cotton mealybug, regarded as an invasive pest worldwide, particularly in the tropics and subtropics. It is one of the major pests of cotton and other commercially important crops. Despite the significant economic losses caused by cotton mealybug the molecular aspects of this insect are under-studied.

**Methods:** In the present study, proteome data of four different developmental stages of cotton mealybug is generated. Differential expression of proteins (DEPs) was studied among six different groups of which, maximum DEPs (550 up-regulated and 1118 down-regulated) were obtained when the quantifiable proteins of Egg+first nymphal were compared with second nymphal instar (FC ≥ 2, P < 0.05). From this proteomics data fifteen potential target genes were predicted for insect pest management. Further, these fifteen genes were explored and evaluated the for RNAi based pest control and optimisation of dsRNA delivery system in cotton mealybug. The analysis of transient expression of target genes was performed.

**Results:** The results signified that dsRNA of *Ferritin* caused ∼69% mortality hence, could be exploited as a promising candidate gene to design a sustainable method for cotton mealybug management.

**Conclusion:** This study provides an urgently required, alternate green control strategy based on proteomics to identify potential RNAi targets for pest management.

## Introduction

*Phenacoccus solenopsis* Tinsley (Hemiptera: Pseudococcidae), commonly known as cotton mealybug, are small-sized, sap-sucking insect pest species that severely impair plant growth and physiology (Arya *et al*., 2017). Additionally, this pest secretes a large amount of honeydew, thus attracting ants, which further leads to fungal infection on the infested plants. The infested plants can show general symptoms of distorted body parts and stunted growth, compromising the yield, and making the cotton mealybug a major pest of fiscal/economic repercussions. It is regarded as an invasive pest worldwide, particularly in the tropics and subtropics. Over 166 plant genera (yielding various industrial and fiber crops) from 51 distinct families are infested by mealybug (Nagrare *et al.,*2011). Owing to the increase in international trade and loopholes in the international quarantine system of quarantine regulations, *P. solenopsis* infestation has spread to all the major cotton producing countries. By the year 2006, in India, mealybug infection was established and reported in all of the nine cotton-growing Indian states (Haryana, Punjab, Rajasthan, Gujarat, Andhra Pradesh, Madhya Pradesh, Maharashtra, Tamil Nadu, and Karnataka) (Dharajyoti *et al*., 2008). In Gujrat, *P. solenopsis* potentially reduced cotton production by 40–50% in 2006. In the same year, in Bhatinda (India), over 2000 acres of cotton production (of which over 100 acres was Bt Cotton), was affected by cotton mealybug infestation. In north India alone, the total damage in 2007 was estimated to be between 400,000 USD and 500,000 USD (Goswami, B., 2007). In both India and Pakistan, this situation resulted in an increase in insecticidal application cost by 250–375 USD per acre. According to a report by the Centre for Agro-Informatics Research (CAIR), Pakistan, the mealybug decimated 0.2 million bales of cotton (170 kg lint per bale) across the country in the year 2007. The report emphasized on the development of preventive measures for mealybug population control. Otherwise, the insect could potentially cause an epidemic in cotton-growing regions over the coming years.

Presently, majority of the control methods are chemical pesticide-based, that are ineffective against the sap-sucking category of insect pests as the contact area (probing area) is much less. Because of being a prominent pest with significant host plants and a high reproduction rate, cotton mealybug spreads quickly and becomes invasive. To stop this spread a highly effective pest management approach is required. Even after the availability of toxins like δ-endotoxins isolated from *Bacillus thuringiensis,* the possibility of applying a transgenic approach for pest control seems ineffective against this sap-sucking insect (Chougule *et al*., 2012). Moreover, limited number of macromolecules are known to protect transgenic crops from mealybugs. Hence, there is an urgent need of devising alternative environment-friendly techniques for *P. solenopsis* management. RNA interference (RNAi) is a natural process in which small RNA molecules regulate gene expression, via sequence specific degradation of target mRNA or by altering the translation process (Mannsperger *et al*., 2010). RNAi has been investigated as a viable method for controlling various insect pest population. RNAi-based gene expression regulation for pest control requires the selection of potential genes and the functional knowledge of the selected genes. The study of insect omics has been demonstrated to be a promising strategy for identifying and selecting novel and species-specific target molecules. The data acquired from the transcriptome and proteome together provide invaluable resources for a systematic investigation of any organism (Ma *et al*., 2014). After identifying a specific target, a disruptive agent can be developed and utilized as a bio-rational pesticide. These disruptive agents, distinguished by their increased selectivity and ecologically friendly nature, are known to interfere with physiological processes unique to a group of insects. This makes them suitable biopesticides (Cusson *et al*., 2008). However, in the cotton mealybug deficiency of proteome sequence, information severely hinders gene discovery, functional validation, expression profiling, and, ultimately, a pest management approach. Advances in mass spectrometry-based proteomics have made the investigation of complex organisms a lot easier (Hewes *et al*., 2001; Tamhane *et al*., 2021). Proteomics-based analysis has considerably enhanced our knowledge of the molecular mechanisms underlying insect metamorphosis, diapause, embryogenesis, vitellogenesis, and their control (Zhnag *et al*., 2001; Zhai *et al*., 2013; Sehrawat *et al*., 2014). A better perspective on signaling, regulatory, and metabolic networks driving insect-specific processes would be possible by integrating proteomics findings with the other molecular approaches. The above information served as an inspiration for this study. The establishment of protein isolation protocol for cotton mealybug, proposing the proteomics data of all four developmental stages and identification of promising candidate genes for pest management of cotton mealybug and related insects are the main foci of this study. Here, the author has combined and compared the proteomics data with the transcriptomics (previously reported by their research group) of all four developmental stages of cotton mealybug. Further, potential target genes were selected from the omics data for RNAi-based gene expression regulation of the target pest. This study concludes that down-regulation of iron storage, transportation, and iron homeostasis maintaining protein *Ferritin* can be used for controlling cotton mealybug population. This is the pioneer study proposing the protein isolation protocol for cotton mealybug, proteomics data of all four developmental stages. Also, this is the first report targeting *Ferritin* genes for insect pest management in the family Pseudococcidae.

## 2. Materials & Methods

### 2.1 Insect culture

*P. solenopsis* population was maintained at CSIR-National Botanical Research Institute (Lucknow, India) from a colony started in 2015 on cotton (*Gossypium hirsutum*) and grown at 28°C temperature and L14:D10 photoperiod. Climate-controlled growth chamber at 28°C, L14:D10 photoperiod, and 60-70% relative humidity (RH) under laboratory conditions were maintained throughout the experiment (Arya *et al*., 2021).

### 2.2 Sample Preparation

Four different developmental stages (Egg+ first nymphal instar, second nymphal instar, third nymphal instar, and female adult) of *P. solenopsis* were collected for protein isolation. Sample was washed using 1% Triton X-100. This step was repeated until the waxy coating was completely removed. Washing of the insect was repeated using deionised water until no frothing was seen in the tube.

### 2.3 Protein extraction, estimation & digestion

500 mg of insect tissue was finely grinded using liquid nitrogen. The powdered tissue was homogenized in extraction buffer (0.5% SDS, 20 mM EDTA, 10 mM DTT, 100mM Tris-HCl (pH 6.8) and 2 mM PMSF). Incubated at 4°C for 2 hours followed by centrifugation (13,000g, 4°C, 10 mins). The supernatant was transferred to fresh tubes. Cold 10% TCA/Acetone and 5mM β-mercaptoethanol (β-ME) were mixed thoroughly to each tube. This mixture was incubated (at -20 °C for 2 hours), centrifuged (at 13,000 g for 5 mins), and the supernatant was discarded (Cilia *et al*., 2009; Niu *et al*., 2018). The protein pellet was vigorously disrupted using a glass rod, washed with 5mL of 80% ice-cold acetone, and air-dried. The pellet was stored at –80°C until further use. Protein pellet was solubilised in rehydration buffer (7M Urea, 2M Thiourea, 2% CHAPS, 100mM DTT) and dialyzed against 20 mM TrisCl, pH 8.0. Protein estimation was performed using BCA method, spectrophotometrically (Supplementary Figure 1) (He F., 2011). The extracted protein was alkylated using 20 mM IAA, and reduced using 10 mM DTT. These procedures were performed at 25°C for 45 min at 37°C for 1 hour, respectively. Further, trypsin digestion was done by adding 100 mM TEAB (Triethylammonium bicarbonate) (Sigma-Aldrich, Darmstadt, Germany) to the protein sample. This step was repeated till the concentration of urea reached <2 M. Trypsin was added such that protein:trypsin ratio of 50:1 ( w/w) was maintained and the solution was incubated overnight. An additional digestion was done with a protein-trypsin ratio of 100:1 for 4 hours. The tryptic peptides were desalted using a C18 SPE column (Phenomenex, Torrance, CA, USA) and vacuum-dried.

### 2.4 LC-MS/MS analysis

The enriched peptides were dissolved in solvent A (0.1% formic acid) and loaded onto a reversed-phase analytical column. The gradient of solvent B (0.1% formic acid in 98% acetonitrile) increased (from 6 to 23% in 26 min, 23 to 35% in 8 min, shot to 80% in 3 min, and held at 80% for 3 min). The experiment was operated on the Acquity Waters UPLC system at a constant flow rate of 400 nL/min. The peptides were then exposed to an NSI source and were analyzed by tandem mass spectrometry (MS/MS) coupled online to the UPLC supplied with 2.0 kV electrospray voltage. (Skipp, *et al*., 2005, Lagrain, *et al*., 2013, Possenti, *et al*., 2013) (Supplementary Figure 2).

### 2.5 Protein identification

The RAW data files were analysed using the Mascot search engines of Proteome Discoverer (Thermo Scientific, Version 1.2). The MS/MS search criteria were as follows: Mass tolerance of 20 ppm for MS and 0.25 Da for MS/MS mode, trypsin as the enzyme with one missed cleavage allowed, carbamidomethylation of cysteine as static, and methionine oxidation as dynamic modifications, respectively. High-confidence peptides were used for protein identifications by setting a target false discovery rate (FDR) threshold of 1% at the peptide level. Only unique peptides with high confidence were used for protein identifications. Further, for peptide ion accounting informatics, PLGS was used as it mines deeper into UPLC/MSE data. And identifies more peptides/proteins with better statistical rigor and sequence coverage than conventional LC/MS/MS methods. Details are listed in supplementary figure 2.

### 2.6 Validation of proteomic data via transcriptomic data and Bioinformatics analysis

The in-house generated transcriptome data of all four different developmental stages of cotton mealybug was used as a frame for proteome data validation. For this validation, the identified peptides were searched/queried against our published transcriptome data of cotton mealybug using PLGS search engine. Further, with the obtained results, bioinformatics analysis was performed. agriGO v2.0 database (http://systemsbiology.cau.edu.cn/agriGOv2/) was used to perform GO annotations. KAAS (http://www.genome.jp/kaas-bin/ kaas_main, version 2.0) were used to perform KEGG analysis.

### 2.7 Differentially expressed proteins (DEPs)

To analyse the differential expression levels of proteins between the major developmental stages, up and down-regulated proteins were estimated between six different groups of developmental stages of *P. solenopsis*. Peptides with two-fold variation in expression with statistical significance between two or more developmental stages were denoted as differentially expressed.

### 2.8 Functional enrichment

#### (a) GO enrichment analysis

As per the GO annotation, proteins were categorized in three groups based on their involvement in biological processes, cellular components, and molecular functions. The enrichment of differentially abundant proteins was performed for each class. The identified proteins were detected using a double-tailed Fisher’s precision test. GO with a P value < 0.05 is considered significant.

#### (b) Pathway enrichment analysis

According to the KEGG database, the differentially expressed protein was enriched against all identified proteins using a two-tailed Fisher’s exact test. The pathway was regarded as significant with a corrected p-value <0.05. These enriched pathways were hierarchically classified with the help of the KEGG website.

### 2.9 Identification of the potential target genes

Peptides present among all the developmental stages of cotton mealybug were shortlisted, their respective gene sequences were retrieved from the in house generated transcriptome data of *P. solenopsis*. The selected genes were studied for their functional annotation, and those involved in vital processes like development, chitin metabolism, digestion and nutrition, detoxification, immunity, pesticide resistance absorption, hormone homeostasis, energy metabolism, cellular and cytoskeletal components were shortlisted for dsRNA efficacy evaluation. dsRNA fragments were designed for the selected genes. Designing potential dsRNA fragments also included limiting the selection of non-specific target genes using bioinformatics. For this, online available dsRNA check software was utilized and similarities between the non-model and closely related species were removed (Naito *et al*., 2009). NCBI-BLAST (National Centre for Biotechnology Information-Basic Local Alignment Search Tool) was used for nucleotide match search for nearly perfect and short sequences were performed against *P. solenopsis* sequence and with the and the *Homo sapiens* genome. GFP was amplified from the binary vector pCAMBIA1303 and, dsGFP was used as negative control.

### 2.10 Sequence analysis of selected genes

Sequence analysis of the selected genes was done using the blastX tool to blast sequences with a non-redundant database of NCBI to annotate their functions. Further, confirmation of the obtained data was done using the in-house generated transcriptome data. Full length gene sequences of the gene fragments that showed higher mortality reported from previous studies and the new transcripts showing the highest ubiquitous expression were used for further analysis. A list of the selected fifteen genes is given in the supplementary table 3 (sequences provided in supplementary file).

### 2.11 Amplification of target genes and dsRNA preparation

The specific dsRNA sequences of selected fifteen target genes are finalized by dsRNA synthesis using cDNA of the third nymphal instar of *P. solenopsis* as a template. The primers designed to amplify target genes had additional T7 promoter sequences (GCGTAATACGACTCACTATAGGGAGA) at the 5’-end and 3’-end. The amplified and purified gene fragments is were further proceeded for dsRNA synthesis expression cassette was created using the MEGA-script kit (Ambion), following manufacturer’s instructions.

### 2.12 Screening of target gene by feeding bioassay

Once the dsRNA expression cassette for all the fifteen target genes of interest and the negative control (GFP) were prepared, these were used for bioassays. Here, for the third nymphal instar we applied two dsRNA delivery approaches. The first approach is soaking method and oral delivery. Experiments were performed as described earlier by Arya *et al*., 2021.

### 2.12a dsRNA feeding bioassays: soaking method

Tobacco leaves were dipped in 200uL solution of 300 ng/μL of dsRNA (and GFP dsRNA as negative control) and nuclease-free water. The experiment was performed in biological triplicates. The process of complete uptake of dsRNA took 4-5 hours. Then third nymphal instar of cotton mealybug (n=10-15) was carefully transferred to these Tobacco leaves. The insects were allowed to feed on dsRNA-soaked Tobacco leaves for 72 hours. A parallel experiment was run for negative control using dsGFP. The experiment was performed in biological duplicates and technical triplicates. Mortality data was evaluated after 72 hours of treatment. SPSS 15.0 software was used for performing statistical analysis. Toxicity assessment of the selected ten dsRNAs against cotton mealybug. In the beginning, a bioassay was performed with dsRNA concentration of 250ng/uL, which was further evaluated for its potency at low concentrations ranging from 40μg dsRNA/mL of diet, 30μg dsRNA/mL diet and 10μg dsRNA/mL diet. Mortality was recorded on the tenth day post treatment. The genes displaying the highest mortality at the lowest time-intervals or low exposure time were further selected for diet-based oral delivery bioassay.

### 2.12b dsRNA feeding bioassays: oral delivery

From the soaking method, ten genes that exhibited mortality of more than 55% were selected for oral delivery-based bioassay. Oral delivery of dsRNA was the second method followed for dsRNA delivery. In the oral delivery method, dsRNA of the selected ten genes were taken in varying concentrations (10, 20, 40 μg dsRNA/mL diet) and mixed with in-house developed and optimized mealybug diet; the third nymphal instars were fed on this mixture (n=10). Insects were transferred to a fresh set of diet (10, 20, 40 μg dsRNA/mL diet) every third day of the experiment. Ten days post-treatment, mortality analysis was done via LC_50_ comparison. The experiment was performed in biological duplicates and technical triplicates. GraphPad Prism software was used for the generation of figures and statistical analysis.

Based on observed mortality from the previous experiments four best-performing dsRNA showing high efficacy at low concentrations, and minimum time in dsRNA uptake were *Ferritin* (Fer), *Cytochrome P450 6a14-like* (P450 6a14), *Cytochrome P450 303a1* (P450 303a1), and *Odorant-binding protein 2 precursor* (OBP) were selected for downstream processing.

### 2.12 c Gene expression evaluation of Fer, P450 6a14, P450 303a1, and OBP

Further, all these four gene were evaluated for their expression. Quantitative RT-PCR analysis was performed to observe gene expression patterns of Fer, P450 6a14, P450 303a1, and OBP at different time intervals of the treatment. GFP was used as a negative control. Comparison was done with the expression levels of untreated insects (control). RNA isolation was done (as described by Arya *et al.,*2021) at zero days (untreated), the eighth day of treatment, and the tenth day of treatment. cDNA was synthesised (as described above). Using gene specific primers, quantitative RT-PCR was performed, as previously reported by Arya *et al*., 2018. Primer 3.0 program was used for primer synthesis. Expression level estimation of target genes in the treated insects was done using the 2^-ΔΔCT^ method. Normalization was done with α*-tubulin* and β*-tubulin.* The selection of housekeeping gene for normalization was influenced from the previously published report by Arya *et al*., 2017 on gene expression analysis at different development phases of *P. solenopsis.* The evaluation was done by taking average of three technical replicates per biological duplicate.

### 2.13 Data analysis

All experiments were performed in triplicate. Graph Pad Prism 5.0 software was used for statistical analysis. The mortality data was recorded and subjected to one-way ANOVA using GraphPad Prism 8.0 (San Diego, California, USA) for each bioassay-response. The standard error mean of the three replicates is represented by an error bar. Asterisk on the graph indicates significance (*P < 0.05, **P < 0.01, ***P < 0.001), among different treatments. Student’s t-test was used for statistical comparisons between groups, and P < 0.05 was considered statistically significant.

## 3. Results

### 3.1 LC-MS/MS and identification of differentially expressed proteins

Using nano-LC-MS/MS and bioinformatics analysis (Schmidt *et al*., 2014) the protein profiling of four different developmental stages of cotton mealybug was performed in the present study. The outline of the steps followed is displayed in Figure 1(a). Differential expression of proteins among different samples of developmental stages of cotton mealybug was observed. Nanoscale liquid chromatography coupled to tandem mass spectrometry (nano LC-MS/MS) technology was utilised and quantifiable proteins obtained from each stage are listed in supplementary table 1. For accessing DEPs six different combinations (Egg+1st nymphal instar **Vs** 2^nd^ nymphal instar, Egg+1st nymphal instar **Vs** 3^rd^ nymphal instar, Egg+1^st^ nymphal instar **Vs** Adult female, 2^nd^ nymphal instar **Vs** 3^rd^ nymphal instar, 2^nd^ nymphal instar **Vs** Adult female, 3^rd^ nymphal instar **Vs** Adult female) were studied. The results of DEPs are represented by the bar graph in Figure 1 (b), 1(c), and 1(d). Among the combinations studied for DEPs, maximum DEPs (1668) were obtained when the quantifiable proteins of Egg+ first nymphal of compared with second nymphal instar, of which 550 were up-regulated and 1118 were down-regulated (FC ≥ 2, P < 0.05). Followed by the group in which proteins of Egg+ first nymphal instar were compared with proteins of adult females in which 1643 DEPs were found, of which 546 were up-regulated and 1097 were down-regulated. Further, Egg+ first nymphal instar Vs. Third nymphal instar displayed 1463 DEPs, of which 523 were up-regulated and 940 were down-regulated. When third nymphal instar Vs Adult female were studied, 534 DEPs were there (149 up-regulated and 385 down-regulated). On comparing the quantifiable proteins of the second nymphal instar Vs third nymphal instar, 202 DEPs were acquired (70 up-regulated and 132 down-regulated). Whereas the least number of DEPs were found in second nymphal instar Vs Adult females, of which 70 were up-regulated, and 40 were down-regulated. The data of this experiment can further be used for exploring the larval-larval, larval-pupal, and pupal-adult molting-related protein expression. From here, we can propose the potential pathways that can be regulated to alter the average growth and development of the insect. Also, the stage-specific molecular pathways can be predicted. As the maximum DEPs were attained by comparing the quantifiable proteins of Egg+ first nymphal and second nymphal instar, it can be proposed that the transition from Egg+ first nymphal to second nymphal instar involves a drastic change at the gene expression level. The in-house generated and published transcriptome data also supported this proposal, as in the transcriptome data, maximum DEGs were obtained Egg+ first nymphal and second nymphal instar (10059 up-regulated and 21581down-regulated) followed by second nymphal instar Vs. third nymphal instar (4487 up-regulated and 242241 down-regulated) (Arya *et al*., 2018).

**Figure 1(a):**
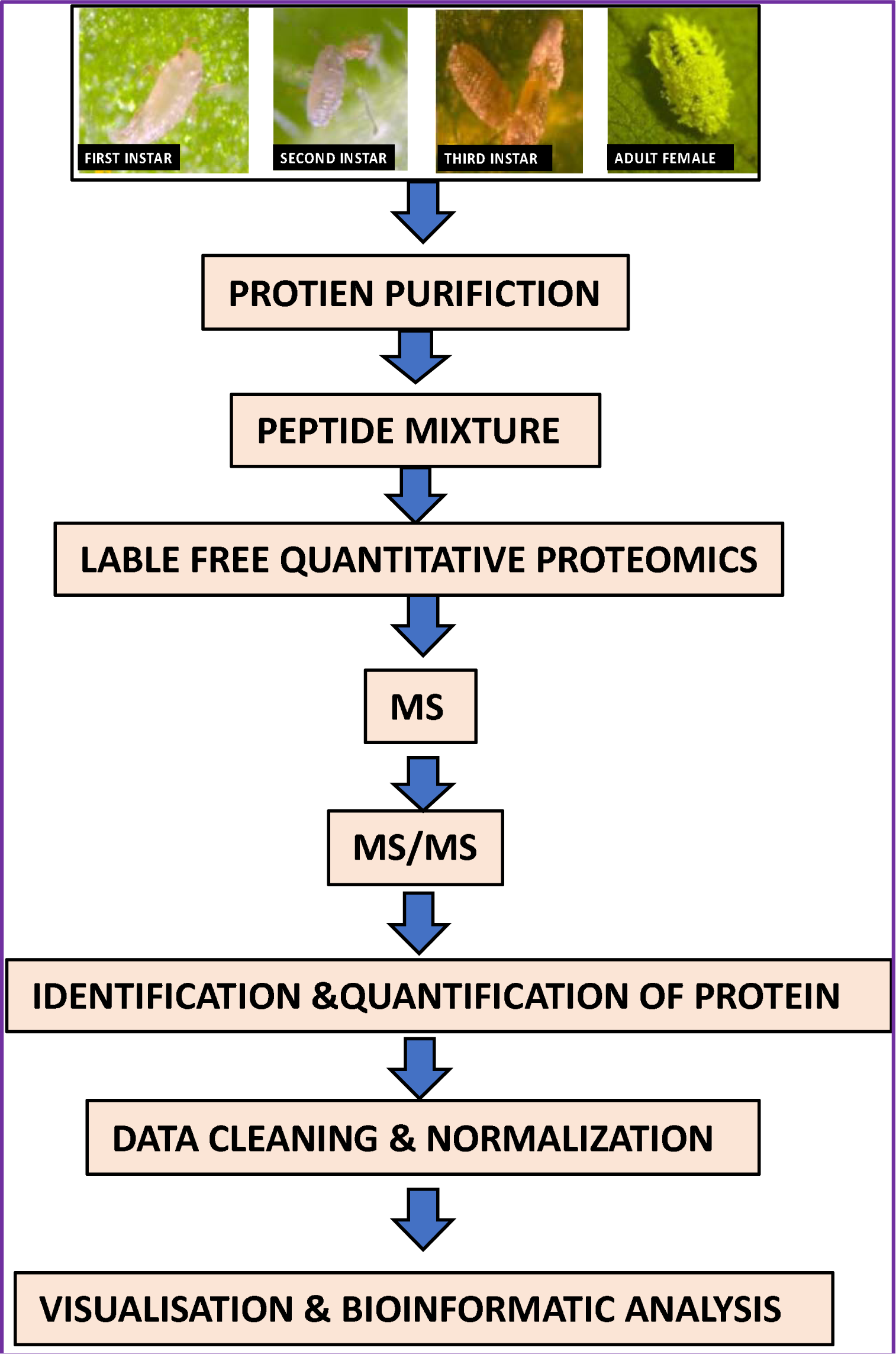
Outline exhibiting steps followed during proteome analysis using nano-LC-MS/MS.

**Figure 1(b):**
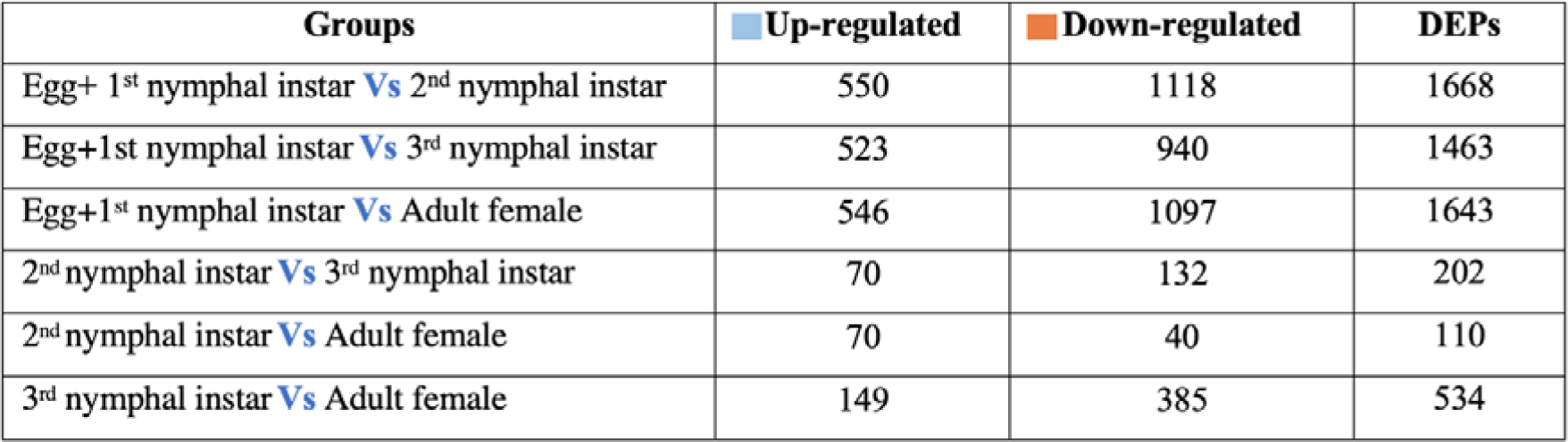
The above table shows the number of unique quantifiable DEPs found in (upregulated are marked in blue and down regulated are marked in orange), in the respective groups among of the four developmental stages in *P. solenopsis*.

**Figure 1(c):**
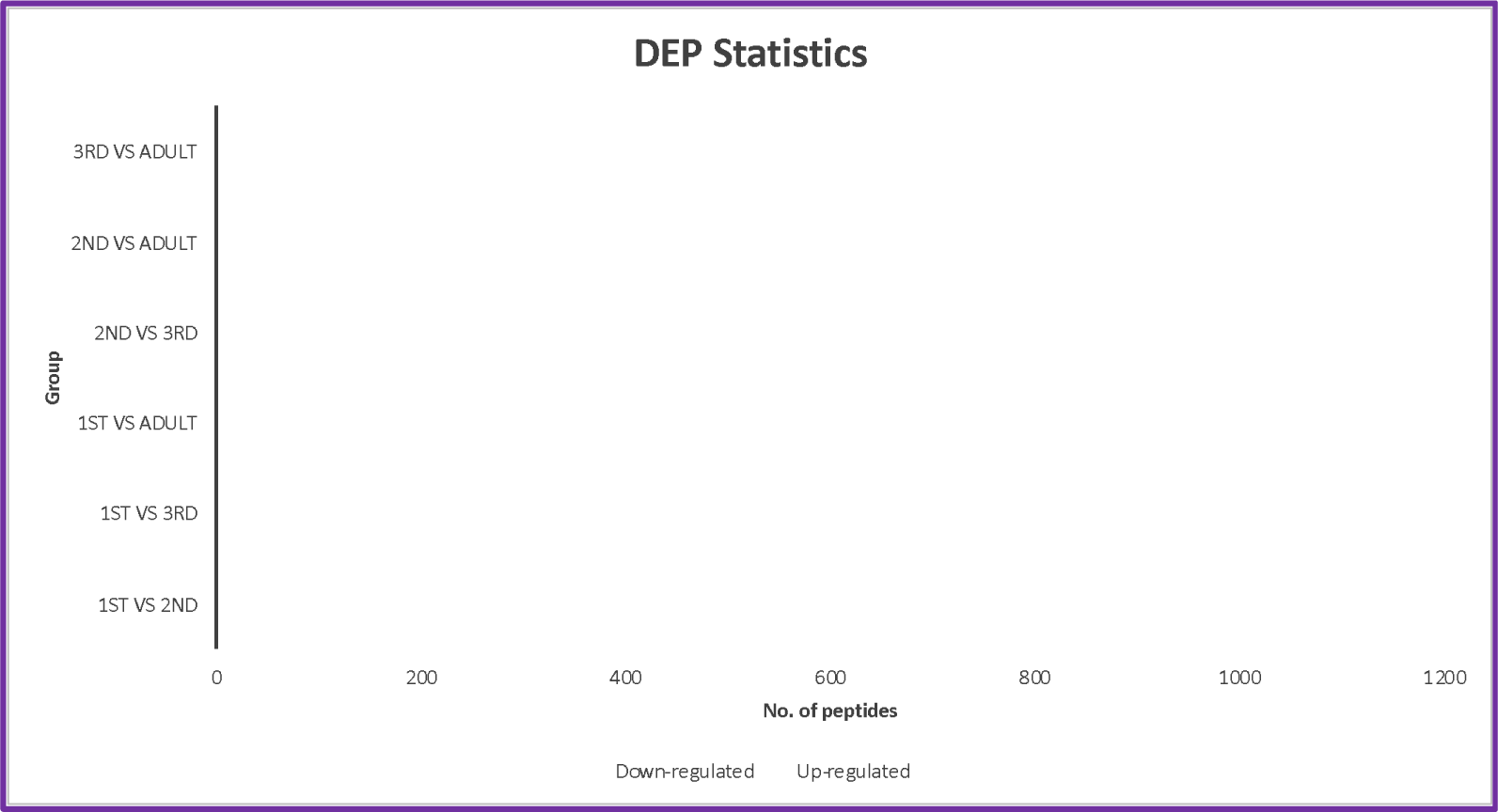
Comparison of the numbers of DEP in different groups of four developmental stages in *P. solenopsis*. Up-regulated peptides are marked in blue, and down-regulated peptides are marked in orange. X-axis represents number of peptides and Y-axis represents different groups. The digits written inside the bars represent the exact count of the unique quantifiable peptides found in that particular category. **NOTE:** The groups studied for DEPs were Egg+ first nymphal instar Vs second nymphal instar (1ST VS 2ND), Egg+ first nymphal instar Vs third nymphal instar (1ST VS 3RD), Egg+ first nymphal instar Vs Adult female (1ST VS ADULT), second nymphal instar Vs third nymphal instar (2ND VS 3RD), second nymphal instar Vs Adult female (2ND VS ADULT), third nymphal instar Vs Adult female (3RD VS ADULT).

**Figure 1(d):**
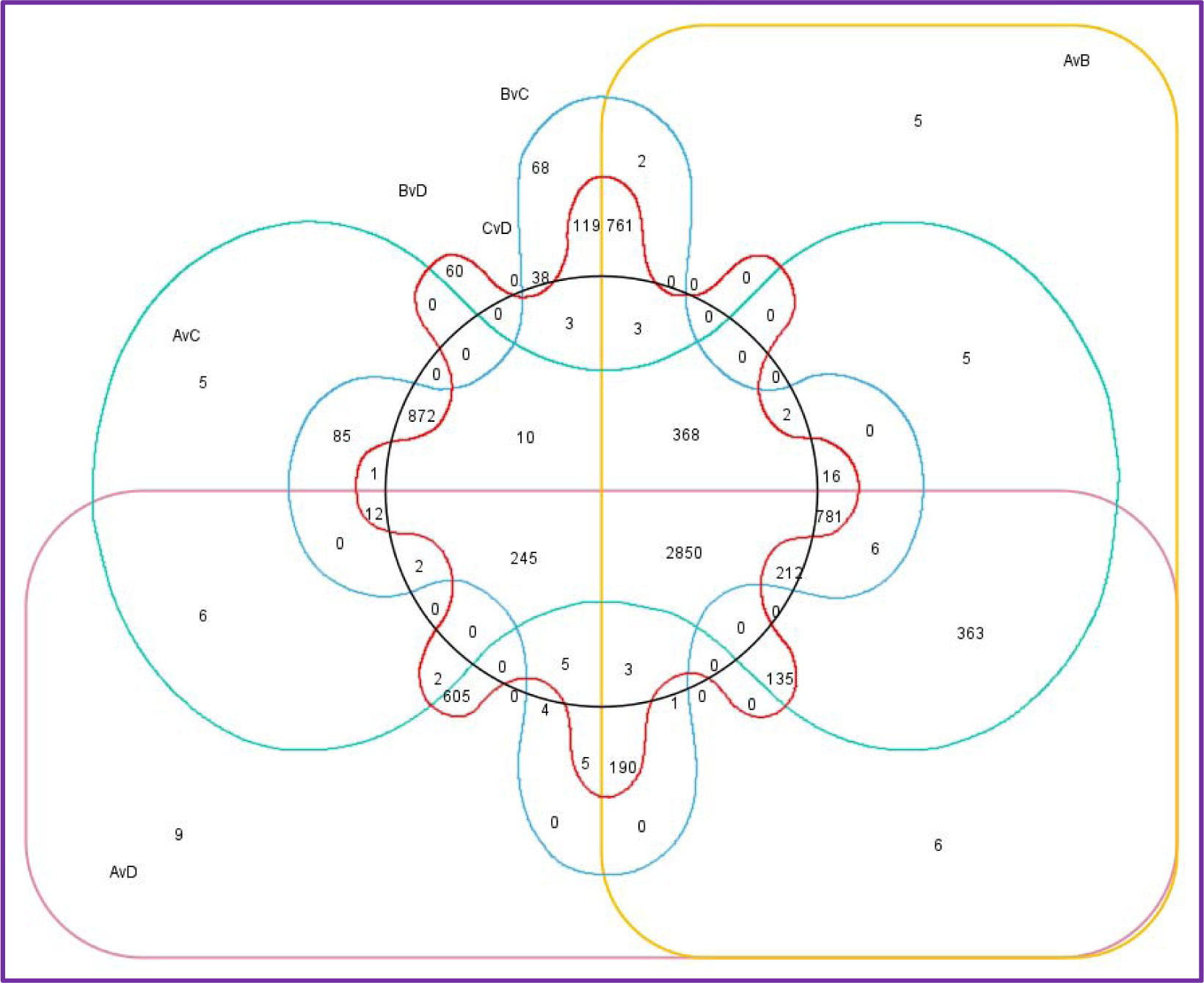
Venn diagrams with number of identified proteins based on nano LC-MS/MS analyses. NOTE: The groups studied for DEPs were denoted as follows: A-Egg+ first nymphal instar, B-Second nymphal instar, C-Third nymphal instar, D-Adult female, AvB-Egg+ first nymphal instar Vs second nymphal instar (1ST VS 2ND), AvC-Egg+ first nymphal instar Vs third nymphal instar (1ST VS 3RD), AvD-Egg+ first nymphal instar Vs Adult female, BvC-Second nymphal instar Vs third nymphal instar (2ND VS 3RD), BvD-Second nymphal instar Vs Adult female (2ND VS ADULT), and CvD-Third nymphal instar Vs Adult female (3RD VS ADULT)

Hence, we can firmly claim that the maximum DEGs and DEPs are found by comparing Egg + first nymphal and second nymphal instar. There is a significant difference in the gene expression and protein expression profile of these two stages, which points towards the involvement of different molting and metamorphosis-related pathways in both developmental stages. Further, the DEPs were categorized based on the function performed. For gene ontology, agriGO v2.0 database was used, and the DEPs were divided into three classes: biological process (BP), cellular component (CC), and molecular function (MF).

A total of, 171 biological processes, 110 molecular functions, and 44 cellular components genes were mapped (Figure 3.8). To understand the pathway information associated with different stages of cotton mealybug KEGG enrichment analysis was performed on the DEPs. The biological pathways of the DEPs were explored using KEGG. In the KEGG assignment, 3,914 quantifiable DEPs were mapped to 307 different pathways. Of these annotated sequences in KEGG, 15.917% of pathways classified were metabolism-related, with most of them involved in carbohydrate metabolism (100 proteins), lipid metabolism (55 proteins), and protein families: metabolism (198), signal transduction (390), cellular processes (258), environmental adaptation (431). A detailed view of the KEGG enrichment is depicted in Supplementary Table 2 and Figure 1 (e) and 1(f).

**Figure 1(e):**
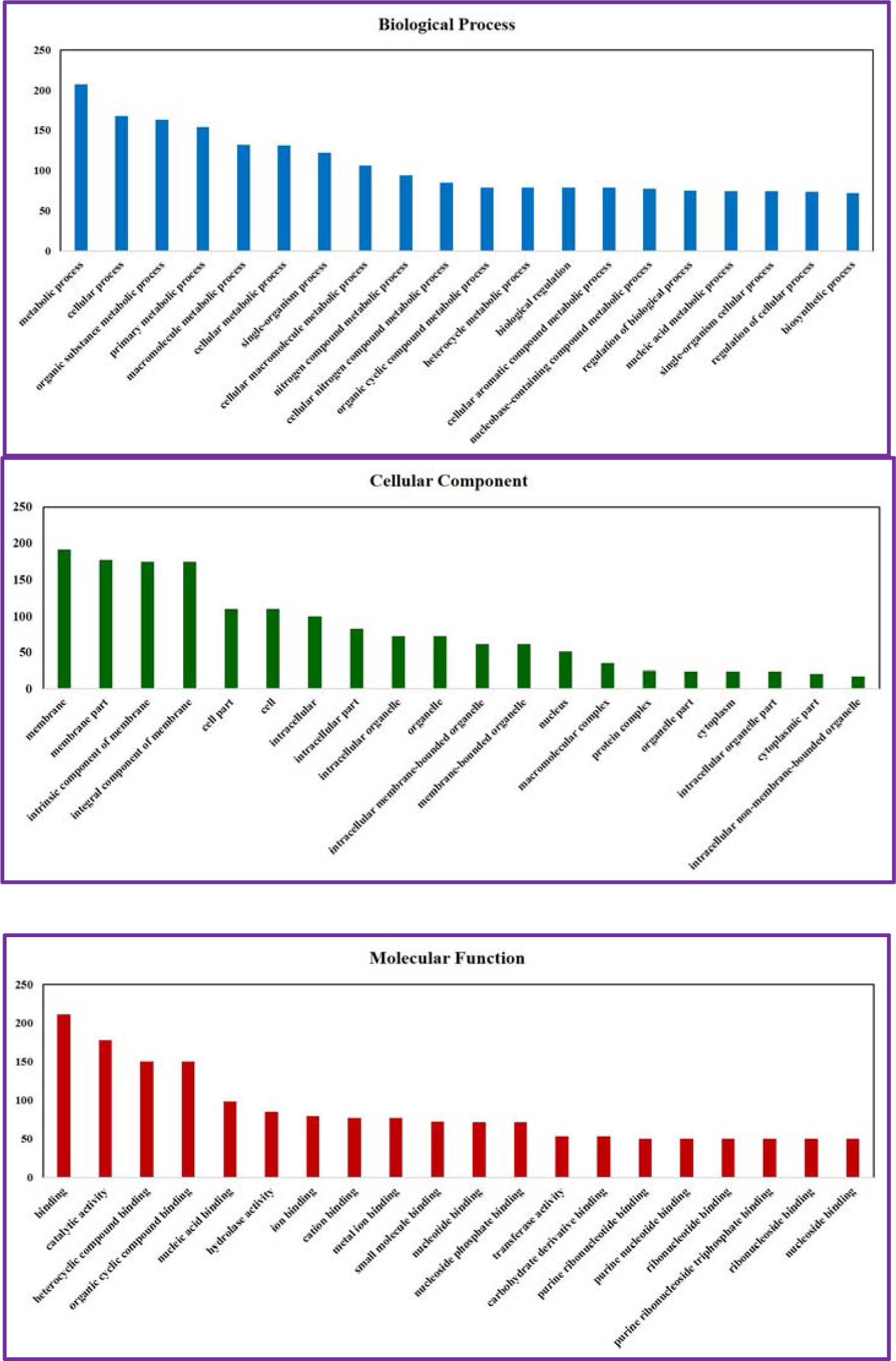
The bar graph is the results of functional classification of DEPs identified in cotton mealybug samples by proteomic analysis. X-axis represents annotation of significant peptides and Y-axis represents number of peptides/proteins.

**Figure 1(f):**
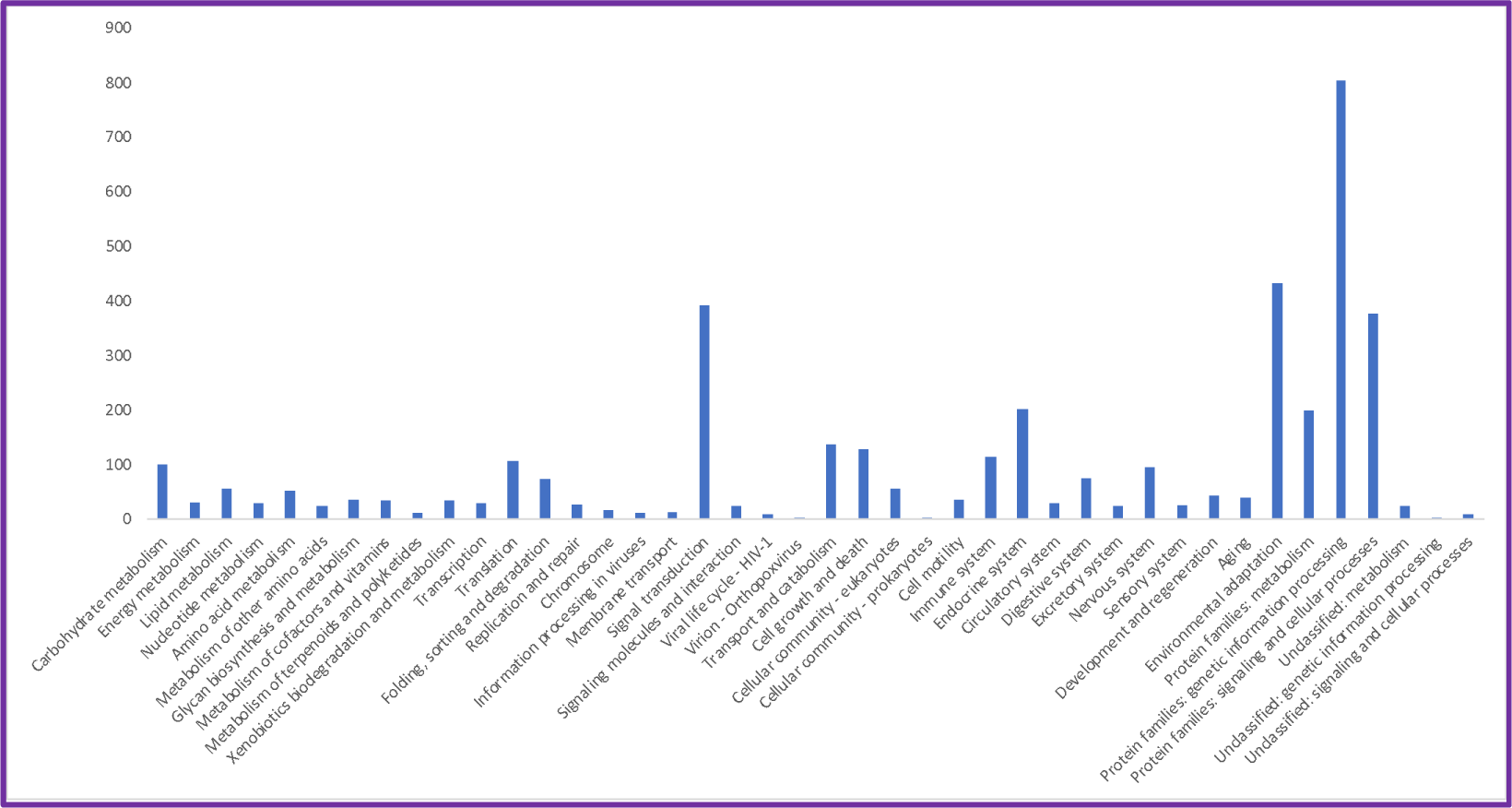
Kyoto Encyclopedia of Genes and Genomes (KEGG) pathway enrichment analysis of DEPs. X-axis represents names of pathways and Y-the axis represents no. of DEPs

### 3.2 Identification of potential RNAi targets

Putative potential genes for inducing mortality in *P. solenopsis* were identified using proteome data. The proteome data was rigorously studied, and the genes present in all four stages were separated. The gene annotation was done using the previously published transcriptomic data of *P. solenopsis* (Arya *et al*., 2018). Further, these genes were matched with the Database of Essential Genes (http://www.essentialgene.org/) with the vital genes of *T. castaneum* (100) and *D. melanogaster* (339) for RNAi-mediated silencing. The genes were filtered as reported by Thakur *et al*., 2016, for selecting RNAi targets in insects. Finally, fifteen genes were selected. Namely, Odorant receptor (ODR), Cytochrome C oxidase (COX), Major Hsp 70(HSP70), Odorant-binding protein 2 precursor (OBP), Importin (IMPRT), Cdc 42 small effector protein (CDC42), probable cytochrome P450 6a14-like (P450 6a14), probable cytochrome P450 303a1-like (P450 303a1), HSP90 cochaperone CDC37-like protein (HSP90), protein Wnt-4 precursor, putative (WNT), chitin binding peritrophin-A, putative (CBP), Ferritin (Fer), chitin-binding domain containing protein (CBD), peroxisomal membrane protein PMP22-like (PMP22), and sugar transporter SWEET1-like (SWEET1). These genes were involved in key processes like energy metabolism, reproduction, detoxification, nutrition absorption, defence and others. GFP was selected as negative control. A detailed list of selected fifteen genes is provided in supplementary table 3. The selected fifteen genes were evaluated for their RNAi efficiency by performing insect bioassays. dsRNA was delivered to target insect, *P. solenopsis* by feeding them on dsRNA (200 ng/mL) soaked Tobacco leaves, mortality was recorded, and graph for percentage mortality was plotted (Figure 2(b)). The maximum mortality was shown by the dsRNAs of *Fer* (75%), *CBD* (74%), *OBP* (70%), *P450 6a14* (67%), and minimum mortality was projected by dsRNAs of *hsp90* (45%), *cdc42*(34%), *pmp22* (32%), *importin* (30%) and *GFP* (21%). Among these target genes, the ten exhibiting maximum mortality (more than 50%) were considered the most potent and carried forward for the downstream evaluation.

**Figure 2(a):**
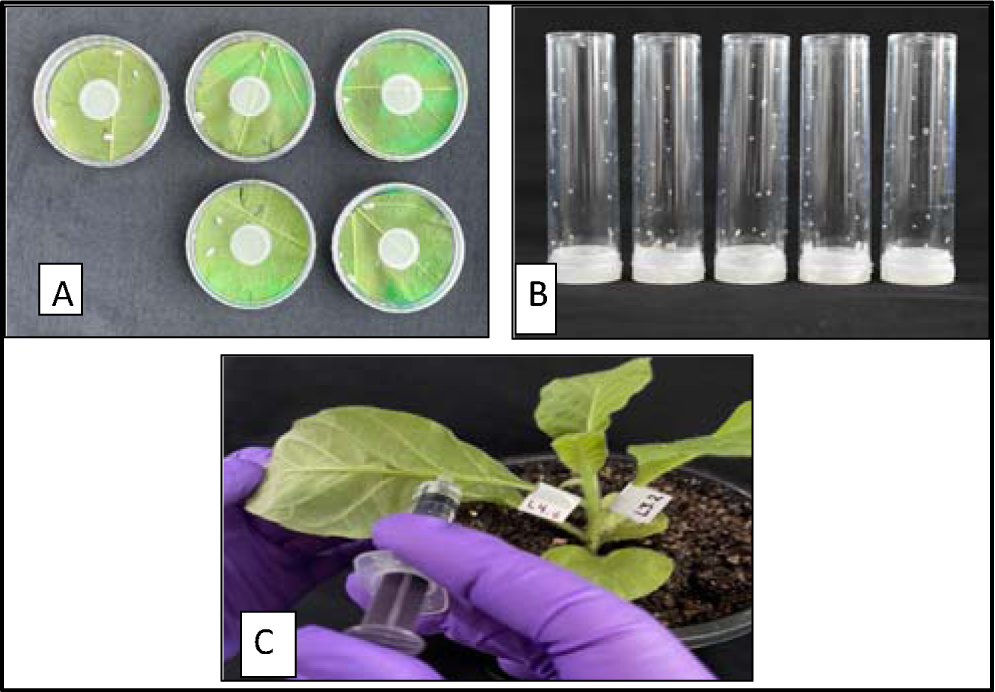
The figure shows the three dsRNA delivery methods applied in this study: A) dsRNA delivery by soaking method, B) dsRNA delivery by artificial diet method, C) dsRNA delivery by agro-infiltration

**Figure 2 (b):**
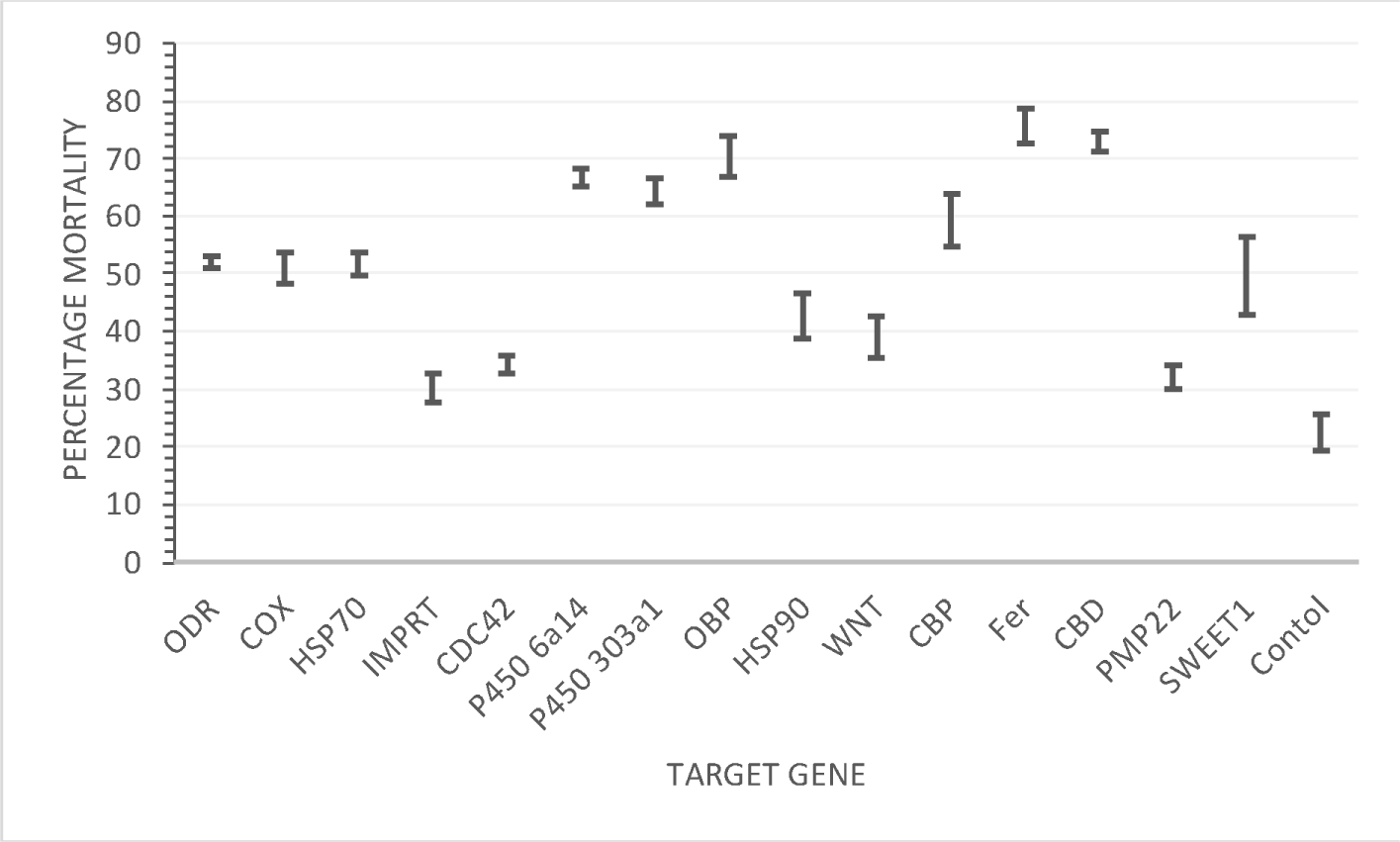
Mortality percentage of fifteen selected genes (GFP as control)targeting the third instar of cotton mealybug after dsRNA (200 ng/mL) delivery by soaking method. In the graph X-axis represents the name of the target genes and Y-axis represents the mortality percentage.

Soaking the Tobacco leaves with dsRNA is an effective method for initial screening but required dsRNA in large quantities. Hence, another bioassay was optimized with an artificial diet. Ten most efficient genes (and GFP), causing mortality >50% were taken in three different concentrations mixed with artificial diet at different concentrations (10, 20, 40 μg dsRNA/mL diet) and mixed with the artificial diet of the insect. Details of these ten selected genes is given in Table 1. The third instar of *P. solenopsis* was allowed to grow on these diets. The data was recorded after 12 hours, 24 hours, and 72 hours of treatment. As the maximum mortality was shown at 72 hours, LC_50_ was calculated for all the dsRNAs after 72 hours of treatment. For the recorded data LC_50_As per the recorded LC_50_ (Table 2), maximum mortality at the lowest concentration was displayed by Fer (11.78 ± 0.00), OBP (17.34 ± 2.28), and P450 6a14 (19.83 ± 0.00). genes reported to show highest LC_50_ were HSP90 (37.54 ± 0.89), WNT (38.8 ± 4.99), and PMP22 (39.78 ± 0.70).

**Table 1:**
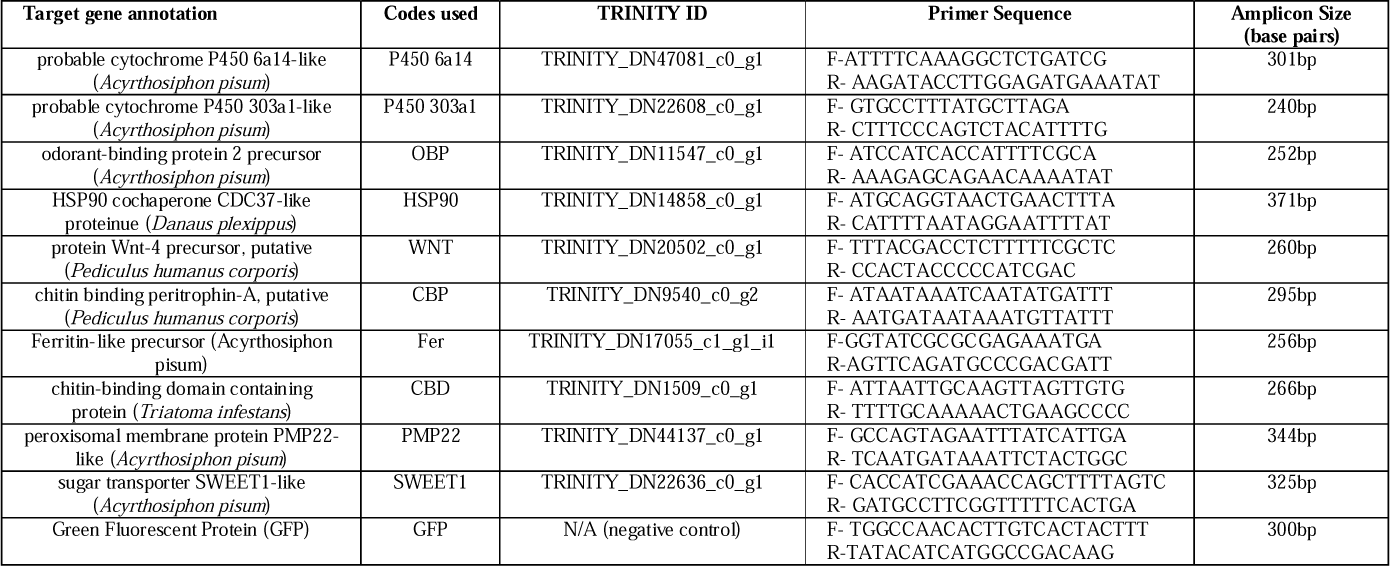
The above table is showing the details of the genes selected for dsRNA feeding bioassays by oral delivery.

**Table 2:**
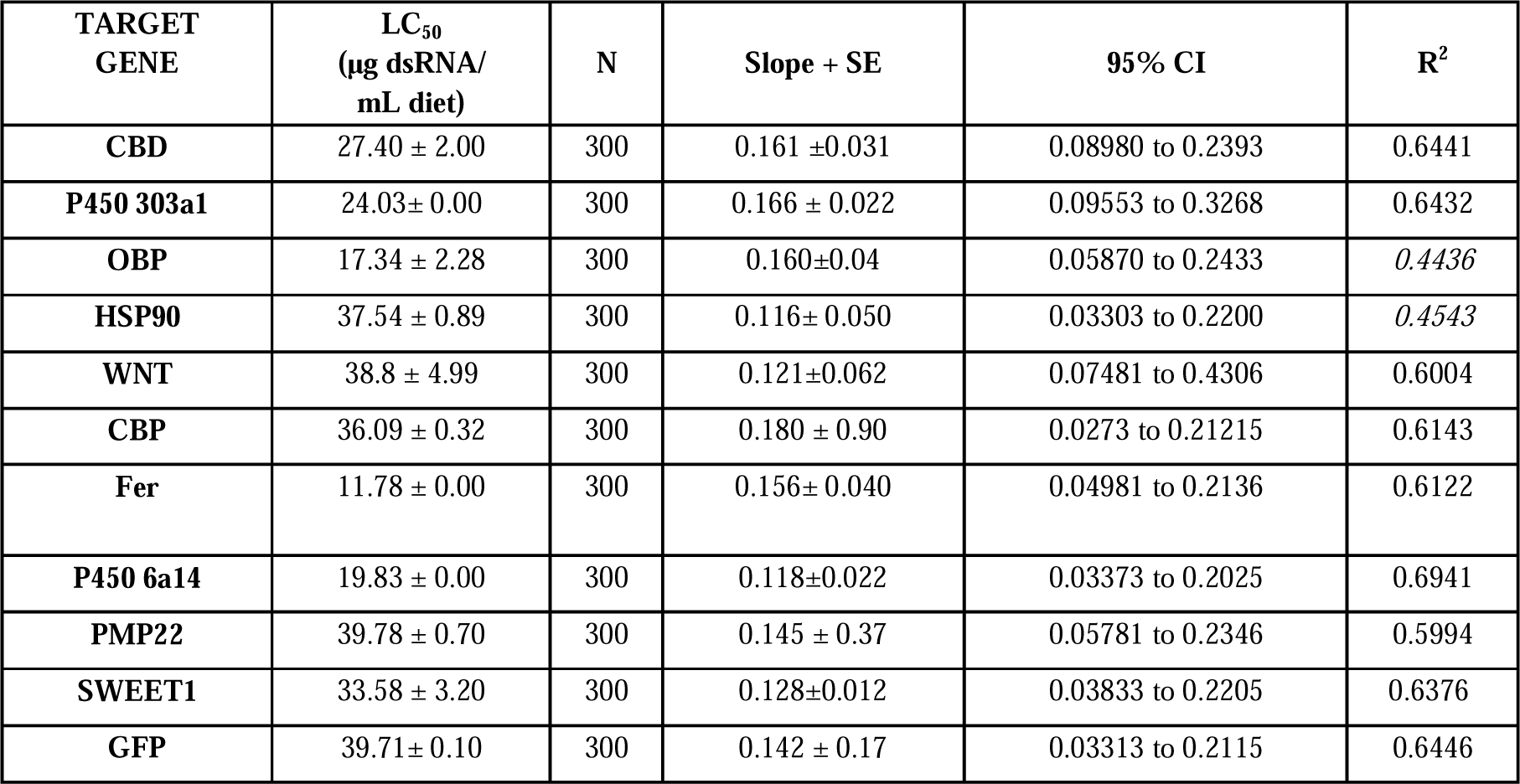
The above table represents LC_50_ (μg dsRNA/mL diet) of dsRNAs after 72 hours of dsRNA feeding treatment with third instar nymphs of cotton mealybug.

Based upon the previous observations, the gene P450 303a1 was expected to show good mortality even at low concentrations, surprisingly, the LC_50_ of P450 303a1 (24.03± 0.00) was higher than P450 6a14 (19.83 ± 0.00) (the other cytochrome variant used in this study). LC_50_ of dsRNA GFP was highest (39.71± 0.00) as per the expectations. The gene GFP is not found in insects; hence, any mortality caused by its dsRNA was minimal and was not due to the downregulation of GFP expression but rather because of unmeasurable and unknown variables having biased influence on the results. Based on LC_50_ of recorded for all the ten target dsRNAs, it was proposed that Fer (11.78 ± 0.00), OBP (17.34 ± 2.28), P450 6a14 (19.83 ± 0.00), and P450 303a1 (24.03± 0.00) were the four highly potent target gene for inducing lethality in insect pests. In addition to this, the efficacy of selected four genes was evaluated via further optimizing, and examining at the lower concentrations to find the best-working dsRNA concentration with the highest impact. To find this, three different concentrations (lower than the previously examined) were used (10μ g dsRNA/mL diet, 30μ g dsRNA/ml diet, and 40μ g dsRNA/mL) under three different time points (after 2 hours, 4 hours and 6 hours of treatment). The graph is showing a mortality percentage graph, and from this data, it was concluded that P450 303a1 did not exhibit up-to-the-mark mortality (Figure 3(a)). Here, the P450 303a1 gene was dropped, and the rest of the three genes proceeded for the downstream analysis.

**Figure 3(a):**
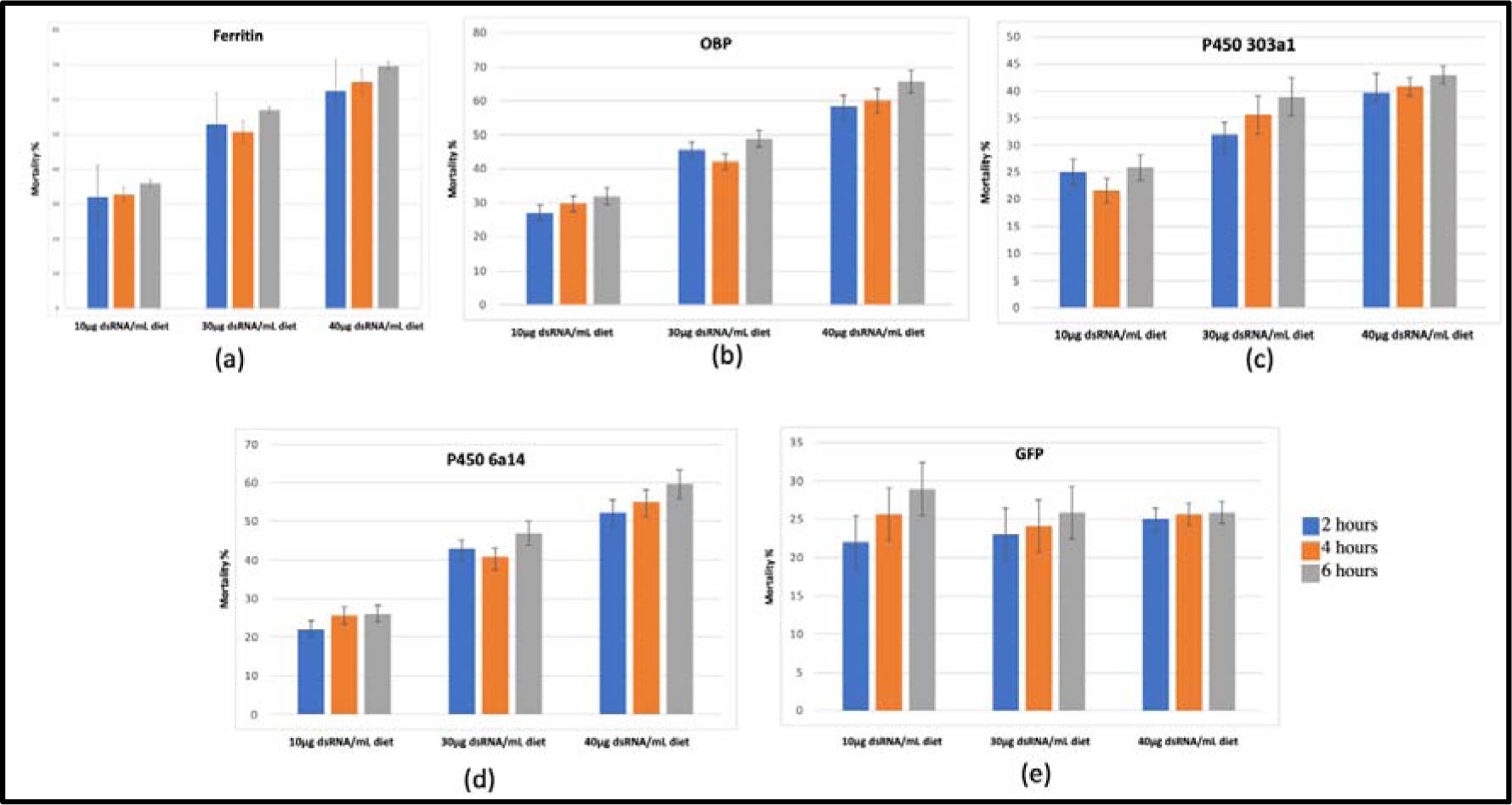
Percentage mortality recorded in *P. solenopsis* on feeding three different concentration ranges (10μ g dsRNA/mL diet, 30μ g dsRNA/ml diets and 40μ g dsRNA/mL) of dsRNAs of five selected genes; *Ferritin* (Fer), *Cytochrome P450 6a14-like* (P450 6a14), *Cytochrome P450 303a1* (P450 303a1), *Odorant-binding protein 2 precursor* (OBP), and dsRNA of GFP (negative control). X-axis represents dsRNA concentration, and Y-axis represents mortality percentage. Note: The experiment was performed in biological triplicates, each with a sample size of 10 individuals of third nymphal instar per treatment group. The figures give the means of these replicates.

### 3.3 Gene expression studies for Fer, P450 6a14, P450 303a1 and, OBP in treated insects

Mortality data and RNAi efficacy data were validated using quantitative RT-PCR. Here, the data was represented in 1/dct form, where the gene expression of individual genes was compared, and relative comparison was not done (Figure 3(b)). dsRNA of GFP was taken as negative control. To strengthen our hypothesis that lethality in the treated insects was stimulated in response to the silencing of the respective gene, RT-qPCR was performed with 10μg dsRNA/mL diet after 8 days of feeding. For Fer, transcript levels were more than 10 folds down-regulated after 8 days of feeding bioassay. The gene expression P450 6a14 gene was 2.75 times down-regulated; for OBP a down-regulation of 2 times was observed when compared to the gene expression of respective genes in the non-treated cotton mealybug population. When the change in gene expression of P450 303a1 was measured, it was found similar in both, the treated and untreated samples with negligible change in gene expression. Most significant down-regulation was obtained in the case of Fer and P450 6a14 followed by the expression of OBP (p < 0.01). The down-regulation was significant (p < 0.01) for Fer, P450 6a14 and P450 303a1 after 8 days of treatment on feeding 10μg dsRNA/mL diet.

**Figure 3(b):**
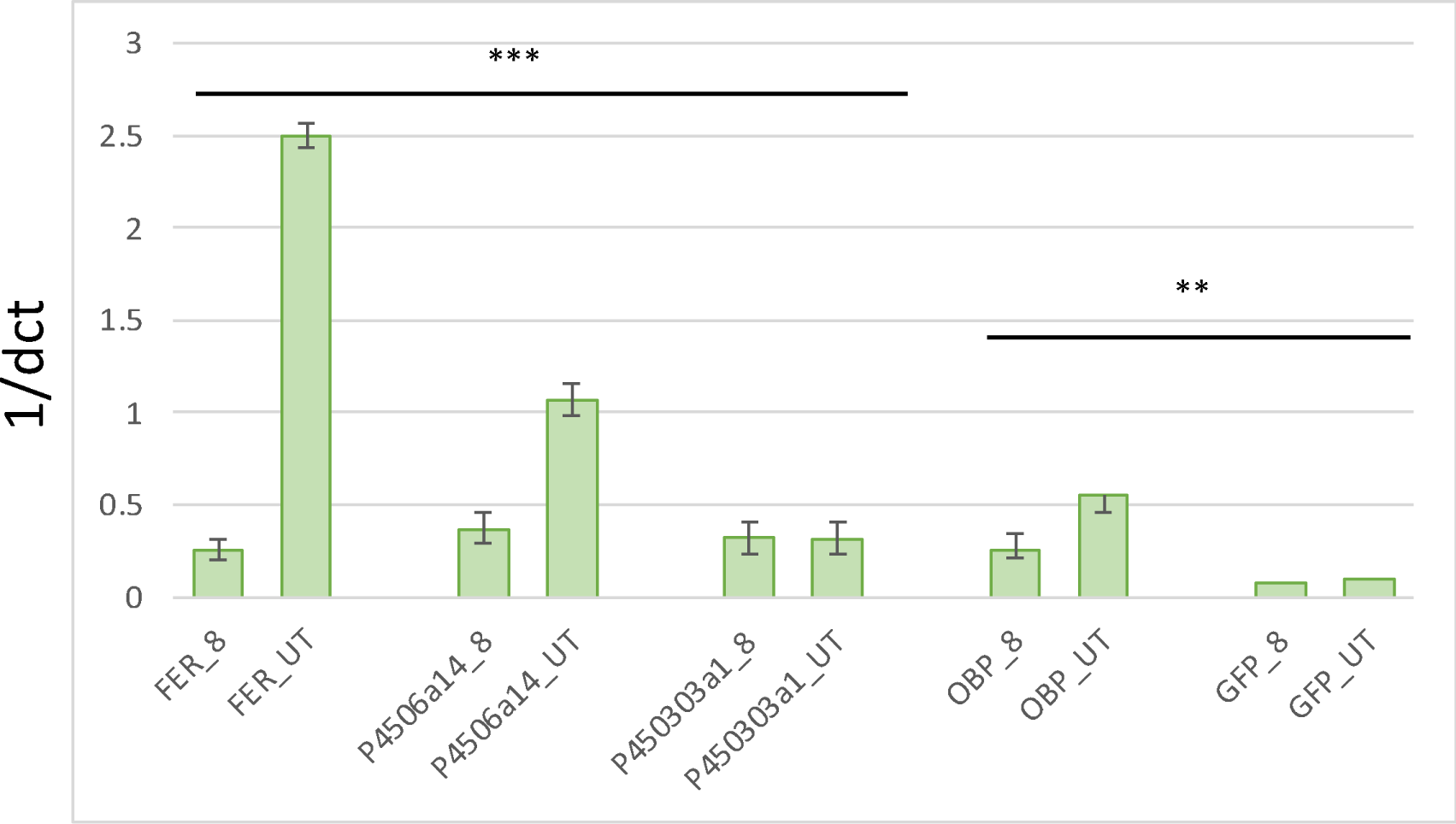
The bar graph shows showing expression of five genes (*Fer, P450 6a14, P450 303a1*, *OBP,* and *GFP*) evaluated through quantitative RT-PCR after oral delivery of dsRNAs via deit-based insect bioassay (10μg dsRNA/mL diet after 8 Days of feeding). Note: The expression of the gene was normalized α*-tubulin* and β*-tubulin,* and GFP dsRNA was taken as negative control (n=3). The graph columns represent the mean ± SE. *P <0:05 and **P <0:01 ***P<0.001 (Student’s t -test). **Note:** Here, Fer_8 represents the expression of the ferritin gene in the insect population (n=5) fed on dsRNA fer + artificial diet (10μg dsRNA/mL diet after 8 days of feeding), and Fer_UT represents an expression of ferritin gene in the insect population (n=5) fed only on artificial die. *P4506a14*_8 represents an expression of *P4506a14*_8 gene in the insect population fed on dsRNA *P4506a14* + artificial diet (10μg dsRNA/mL diet after 8 days of feeding) and P4506a14_UT represents an expression of *p4506a14* gene in the insect population (n=5) fed only on artificial die. P450303a1_8 represents an expression of *p450303a1* gene the insect population fed on dsRNA P450303a1 + artificial diet (10μg dsRNA/mL diet after 8 days of feeding), and P450303a1_UT represents an expression of *p450303a1* gene in the insect population (n=5) fed only on the artificial die. OBPi_8 represents the expression of *OBP* gene in the insect population fed (n=5) fed on dsRNA OBP+ artificial diet (10μg dsRNA/mL diet after 8 days of feeding), and OBP_UT represents an expression of *OBP* gene in the insect population (n=5) fed only on artificial diet. And, GFPi_8 represents the insect population fed on dsRNA GFP + artificial diet (10μg dsRNA/mL diet after 8 days of feeding) and GFP_UT represents expression of *GFP* gene in the insect population (n=5) fed only on artificial die.

### 3.7 Development of gene constructs for the expression of dsRNA in tobacco

From the *in-vitro* bioassay it was inferred that out of the tested fifteen genes *Ferritin* (Fer), *Cytochrome P450 6a14-like* (P450 6a14), and, *Odorant-binding protein 2 precursor* (OBP) exhibited maximum mortality with significant down-regulation of target transcript level at very low concentration. Therefore, these three genes were selected for *in-planta* validation. Plant binary vector pBI101 was used for designing *Ferritin* (Fer), *Cytochrome P450 6a14-like* (P450 6a14), and, *Odorant-binding protein 2 precursor* (OBP), and GFP silencing construct under the control of double enhancer CamV35S promoter as previously reported by our group (Thakur *et al*., 2014). All four (pFERi, pP450 6a14i, pOBPi and, pGFPi) vectors were confirmed by restriction digestion analysis (Figure 4(a)) and figure 4 (b) is showing graphical representation of gene silencing construct maps of all the four selected genes. Individual transformation of all four (pFERi, pP450 6a14i, pOBPi, and, pGFPi) vectors was done in *Agrobacterium tumefaciens* LBA4404 cell by the process of electro-transformation. Further, these transformed *Agrobacterium* cells were finally used for agro-infiltration.

**Figure 4(a):**
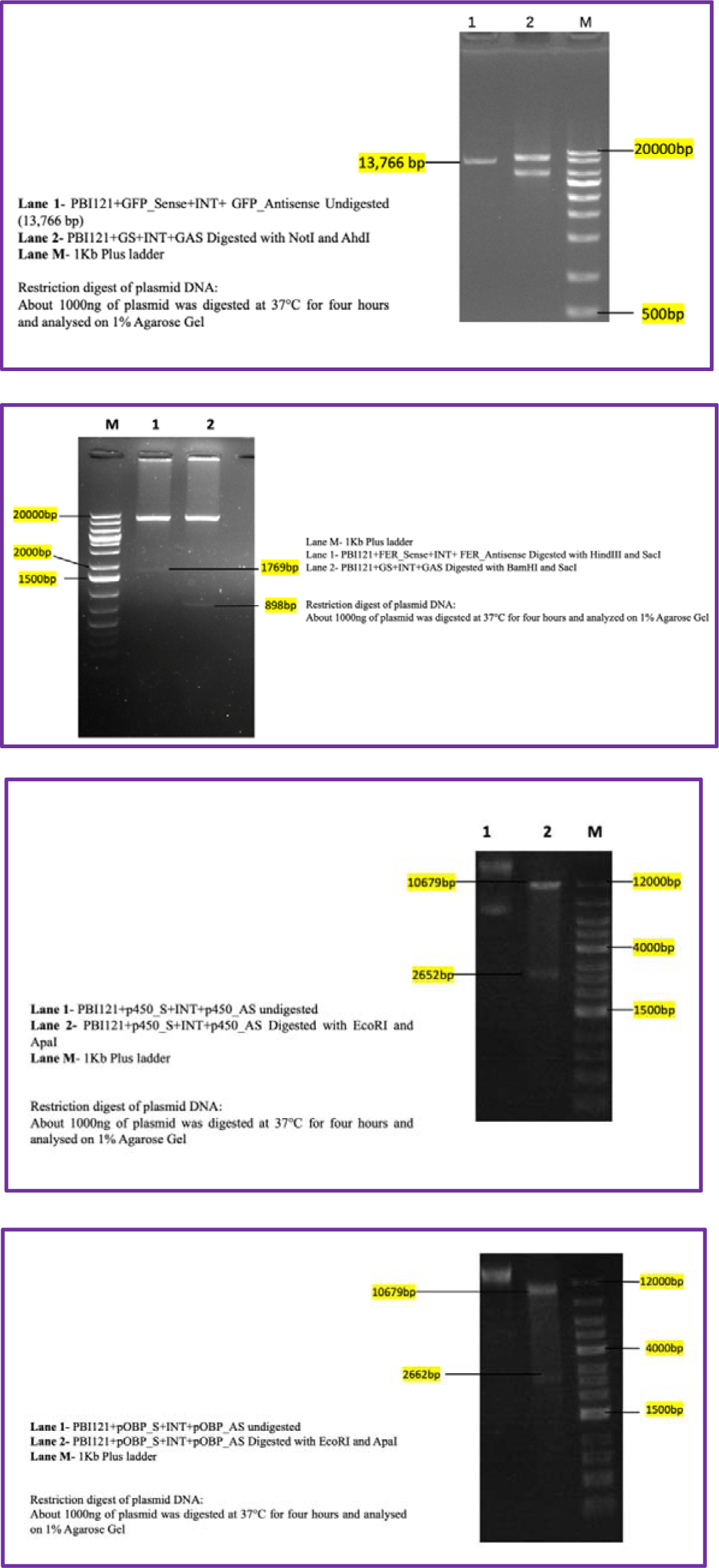
Gel images showing restriction digestion analysis of each individual vector.

**Figure 4(b):**
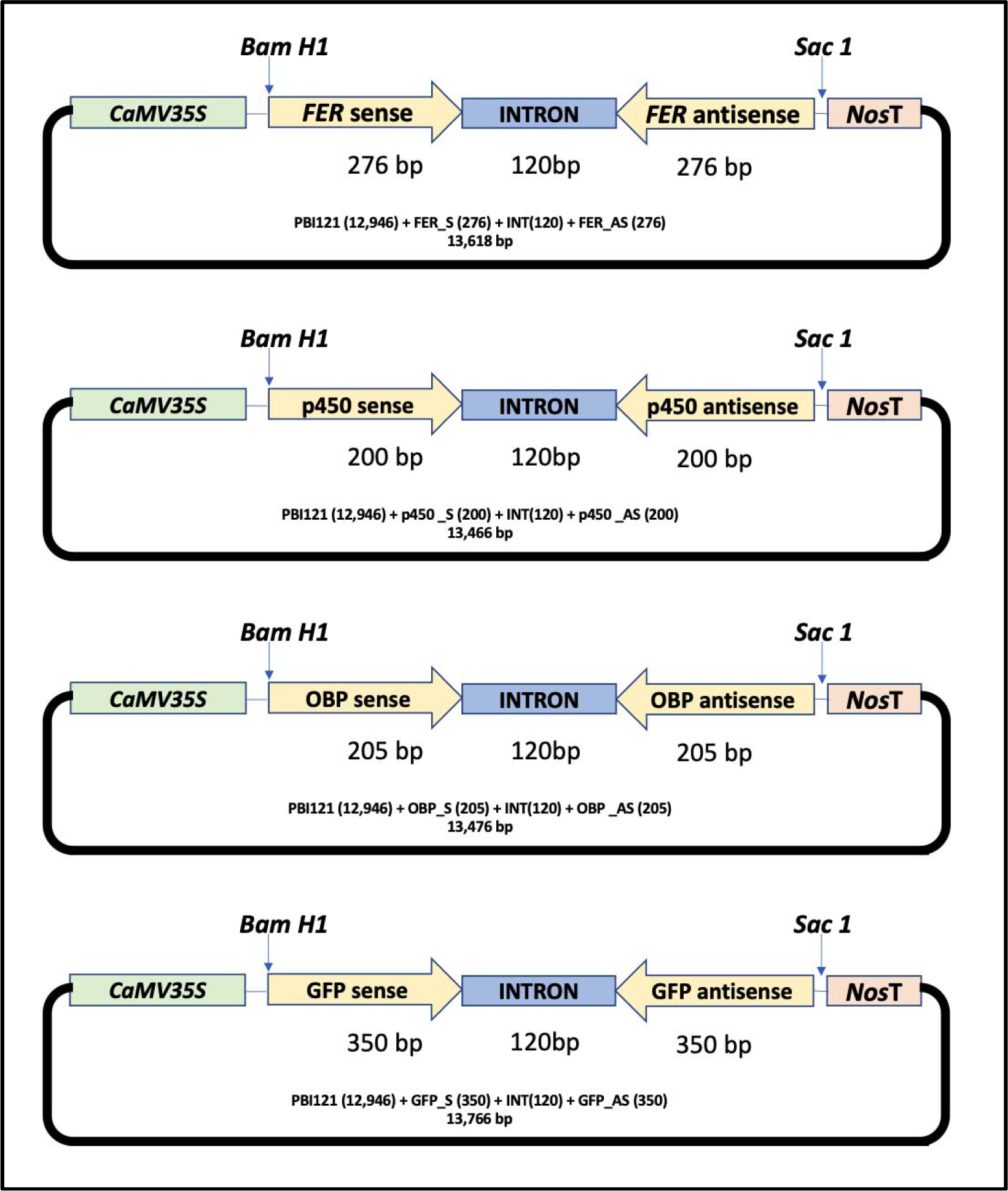
Schematic representation of plant expression Vector pFERi, pp450i, pOBPi and pGFPi.

### 3.4 Effect of RNAi on target gene expression

Tobacco leaves were individually infiltrated with pFERi, pP450 6a14i, pOBPi and, pGFPi vectors. Bioassay was performed with a third instar with two-days-old infected leaves and data was recorded at constant time interval. pGFPi infiltrated tobacco leaves were taken as negative control for insect bioassay experiment. The quantitative RT-PCR results showed significant down regulation in the case of respective genes in case of pFERi fed insects followed by insects fed on pP450 6a14i, and pOBPi agro-infiltrated leaves. The results were similar in case of pGFPi fed insects when compared with non-treated insects (Figure 4(c)). Further, RT-PCR data of expression knockdown was correlated with %mortality data of third instar fed on pFERi, pP450 6a14i, and pOBPi agro-infiltrated leaves showed significant mortality as compared to control (pGFPi agro-infiltrated leaves). Third Instar fed on pFERi displayed maximum mortality of 68.76±3.11 % after ten days of feeding and 62.56±2.86% after 8 days of feeding on agro-infiltrated tobacco leaves. pFERi agro-infiltrated leaves caused 62.56±2.86 % mortality after 8 days of feeding and 68.76±3.11 % mortality after 10 days of feeding. Insects fed on pP450 6a14i agro-infiltrated leaves displayed 55.77±6.86 % and 58.36±5.86 % mortality after 8 and 10 days of feeding, respectively. In the case of pOBPi agro-infiltrated tobacco leaves after 8 days of feeding 52.60±5.86 % mortality was observed and after 10 days of feeding 58.09±6.88 % mortality was observed. However, in the case of negative control, negligible mortality of 10.16±3.60 % was obtained after 8 days of feeding and 15.76±5.86 % mortality after 10 days of feeding (Table 3).

**Figure 4(c):**
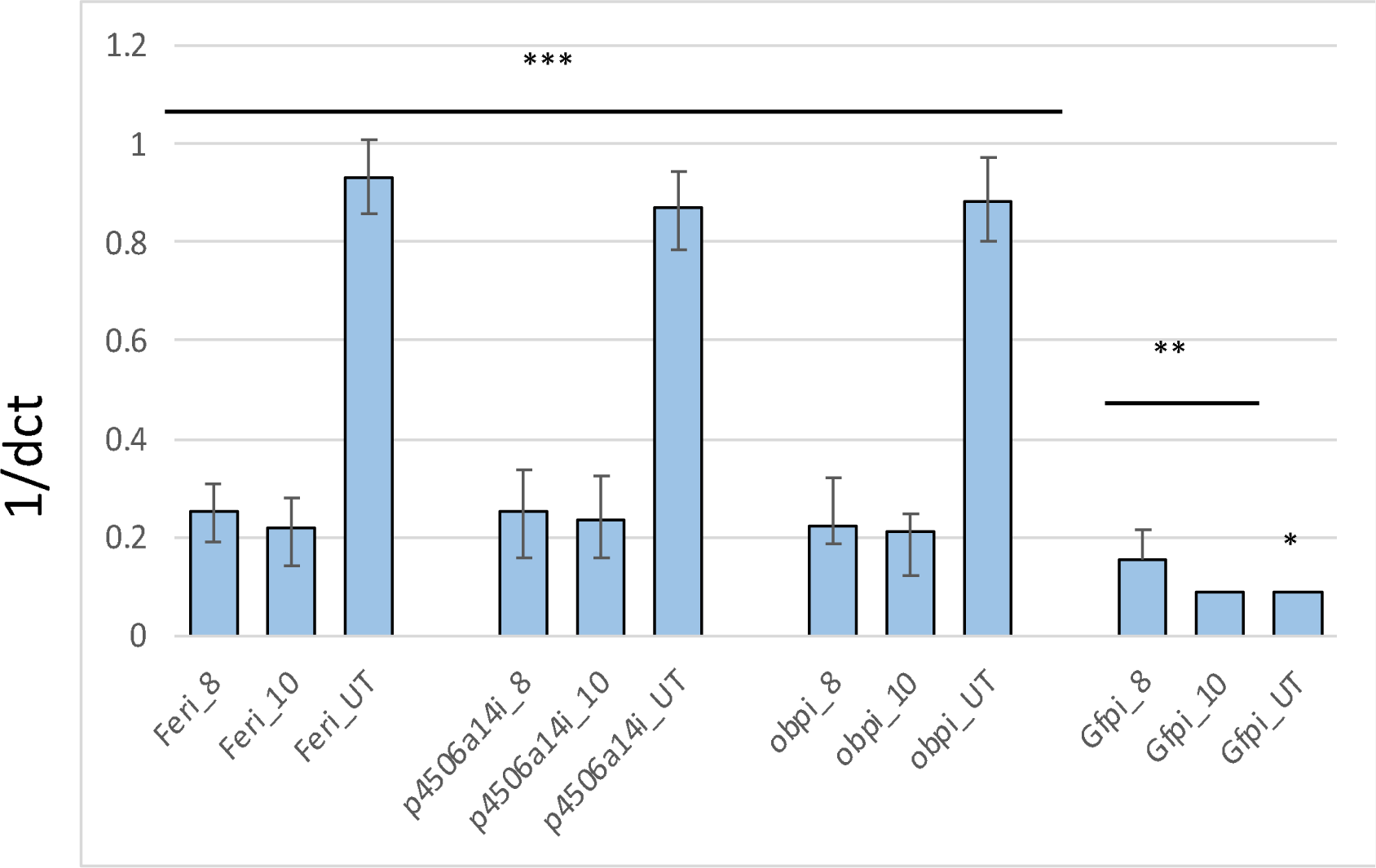
Bar diagram is showing expression of three genes *Ferritin* (Fer), *Cytochrome P450 6a14-like* (P450 6a14), and, *Odorant-binding protein 2 precursor* (OBP) was evaluated through quantitative RT-PCR post-agro-infiltration. The expression of gene was normalized to negative controls that were exposed to pGFPi. The *actin* gene was used as a reference gene (n=3). The graph columns represent the mean ± SE. *P <0:05 and **P <0:01 ***P<0.001 (Student’s t -test). The Y-axis shows 1/dct (relative expression value) and X-axis is showing time interval and treatment. **Note:** Here feri_8 and feri_10 represents the expression of *fer* gene in insect population fed on pFERi infiltrated tobacco leaf for 8 days and 10 days, respectively. P4506a14_8 and P4506a14_10 represents the expression of *P4506a14* gene in insect population insect population fed on pP4506a14i infiltrated tobacco leaf for 8 days and 10 days, respectively. pOBPi_8 and pOBPi_10 represents the expression of *OBP* gene in insect population fed on pOBPi infiltrated tobacco leaf for 8 days and 10 days, respectively. And, pGFPi_8 and pGFPi_10 represents the expression of *gfp* gene in insect population fed on pGFPi infiltrated tobacco leaf for 8 days and 10 days, respectively (used as negative control).Fer_UT is non-treated (control) insect checked for expression levels of FER, P4506a14_UT non-treated (control) insect checked for expression levels of P4506a14, OBP_UT non-treated (control) insect checked for expression levels of OBP and GFP_UT non-treated (control) insect checked for expression levels of P450.

**Table 3:**
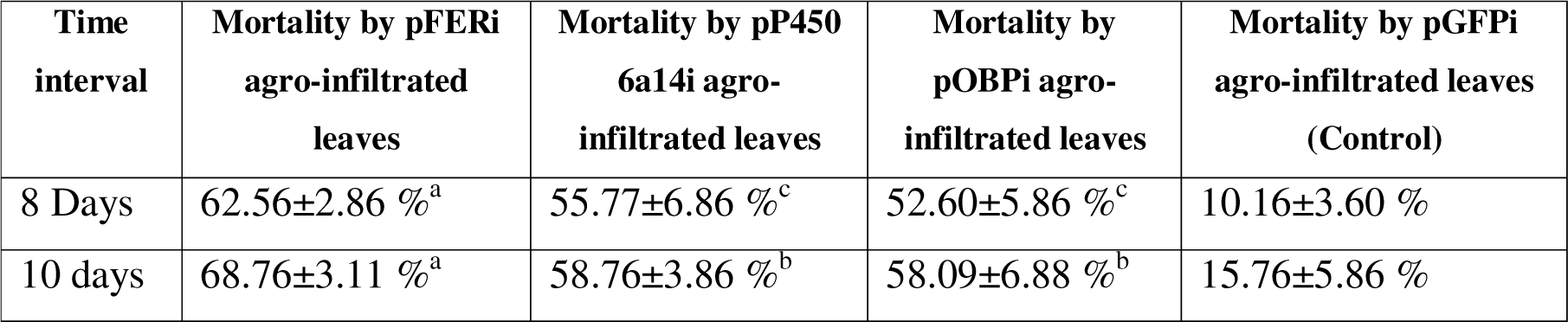
Insect bioassay of the third instar fed on pFERi, pP450 6a14i, pOBPi and, pGFPi infiltrated leaves. Mean ± SE was used for calculating mortality percentage and analysis was done with one-way ANOVA (P<0.05). Means were compared using the DMRT at (P<0.05) (SPSS software); means superscripted with the same letter within a column are not significantly different.

## 4. Discussion

Agricultural limitations must be reduced to meet the needs of the growing population. Pesticides, biocontrol agents, and the introduction of pest resistance in plants have all assisted farmers in enhancing the yield (Lavergne *et al*., 2020). However, it is critical to investigate new methods for combating insect pests in light of recurring reports of field technology failure, such as Bt technology, and the development of resistance against common chemical pesticides. There is a continuing need to seek a better alternative to the approaches for pest management. India is one of the major producers of Cotton, and a significant part of the country’s economy depends on agriculture. Crop growers in India face many challenges, like insect pests. Here, the authors aim to provide an environment-friendly and sustainable option for pest management against cotton mealybug and related pest species.

The author’s research group at CSIR-NBRI, Lucknow (India) and the Department of Botany, University of Lucknow, works on understanding the molecular basis of crop-pest interactions and crop improvement. Both the labs work with pests like *Helicoverpa, Spodoptera, Bemisia tabaci,* and *P. solenopsis,* which are reported as major crop pests of India (Jain *et al*., 2023, Arya *et al*., 2021, Dixit *et al*., 2020). In a previous publication by the same research lab, transcriptome libraries of four different developmental stages of cotton mealybug was generated and information regarding the genomic resources of this pest was provided (Arya *et al*., 2018). Even though transcriptome offers a detailed overview of global gene expression, proteomic data provides a comprehensive insight into the protein profile of an organism hence, used as a complementary approach for better understanding molecular interactions. This information led to the origin of this research. Proteome of four different developmental stages of *P. solenopsis* was performed and validated using the in-house produced transcriptome libraries. Further, the vital genes were identified and evaluated for RNAi efficacy. Ultimately, this study provided three promising gene targets for gene silencing-based control of *P. solenopsis* and related pest species.

This report aims to provide a sustainable and alternative control technique for regulating the cotton mealybug population. The lack of omics-related information on the target pest was one of the most significant challenges faced during this investigation. To achieve this through biotechnological interventions, nano-LCMS was performed, and proteome data for four developmental stages of *P. solenopsis* was generated. Our quantitative proteome study shows, the in-house generated proteome database was studied to select putative target genes for RNAi-based control of *P. solenopsis* and related species. RNAi technology was applied to achieve this goal. The second major challenge was the lack of studies on the possible and effective ways to deliver dsRNA molecules to induce gene silencing. Different dsRNA delivery methods were optimized and evaluated for cotton mealybug mortality during this study. Ultimately, fifteen genes were finalized for their RNAi efficacy evaluation.

Iron is essential for cell viability as it is a co-factor in many cellular processes (Andrews, N. C., 2008). Ferritin has a well reported role in iron absorption, metabolism and homeostasis. Altering the iron metabolism in mosquito can be utilised as a control strategy for regulating fecundity, populations, and transmission rates of mosquito and thereby the vector-borne diseases (Geiser *et al*., 2019). In Whitefly, salivary ferritin is reported to have role in host exploitation by suppressing the plant defence (Su *et al*., 2019). Additionally, insect iron metabolism is significantly different from mammalian ferritins. Insect ferritins are heavier mass, their subunits are usually glycosylated and they composed of a signal peptide. The crystal structure of insect ferritin exhibits tetrahedral symmetry and that of mammalian ferritin shows an octahedral symmetry. The primary association of insect ferritins is with the vacuolar system and iron transport. On the contrary the mammalian ferritins, are cytoplasmic are involved in iron storage (Pham & Winzerling.,2010). Hence, targeting ferritin for pest population management could be a promising approach with minimal non-target effects (especially in mammals).

From the present study it is inferred that RNAi-based gene expression downregulation of a vital gene such as ferritin can be used as a ensuring strategy for insect pest control. This report proposes a sustainable pest management strategy for controlling the invasive pest of cotton, *P. solenopsis.* The data and information from this study can further be utilized to deliver substantial outcomes in the field of integrated pest-management strategies.

## Conclusion

This report aims to provide a sustainable and alternative control technique for regulating the cotton mealybug population. The lack of omics-related information on the target pest was one of the most significant challenges faced during this investigation. To achieve this through biotechnological interventions, nano-LCMS was performed, and proteome data for four developmental stages of *P. solenopsis* was generated. Furthermore, the *in-house* generated proteome database was studied to select putative target genes for RNAi-based control of *P. solenopsis* and related species. RNAi technology was applied to achieve this goal. The second major challenge was the lack of studies on the possible and effective ways to deliver dsRNA molecules to induce gene silencing. Different dsRNA delivery methods were optimized and evaluated for cotton mealybug during this study (Figure 4 (a)). Ultimately, fifteen genes were finalized for their RNAi efficacy evaluation. From the present study, an inference can be made that RNAi-based downregulation of gene expression can be used as a promising strategy for insect pest control. This report proposes a sustainable pest management strategy for controlling the invasive pest of cotton, *P. solenopsis*.

**Institutional Manuscript ID**-CSIR-NBRI_MS/2024/02/13

---

**Supplementary Figure 1a:**
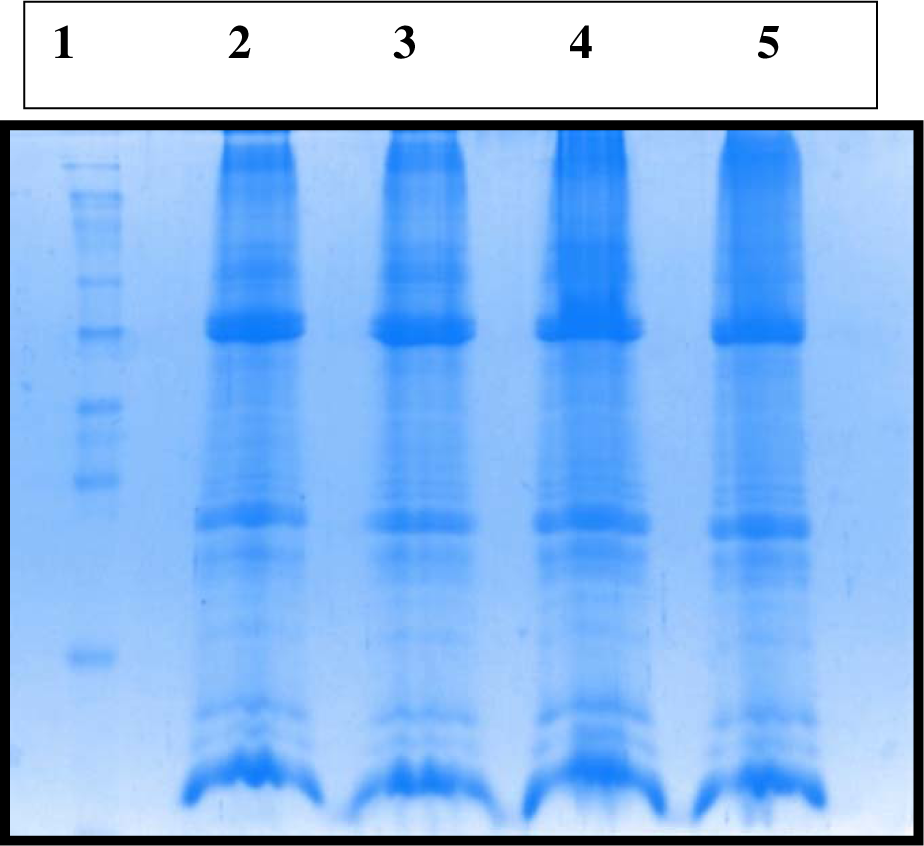

**Supplementary Figure 1b:**
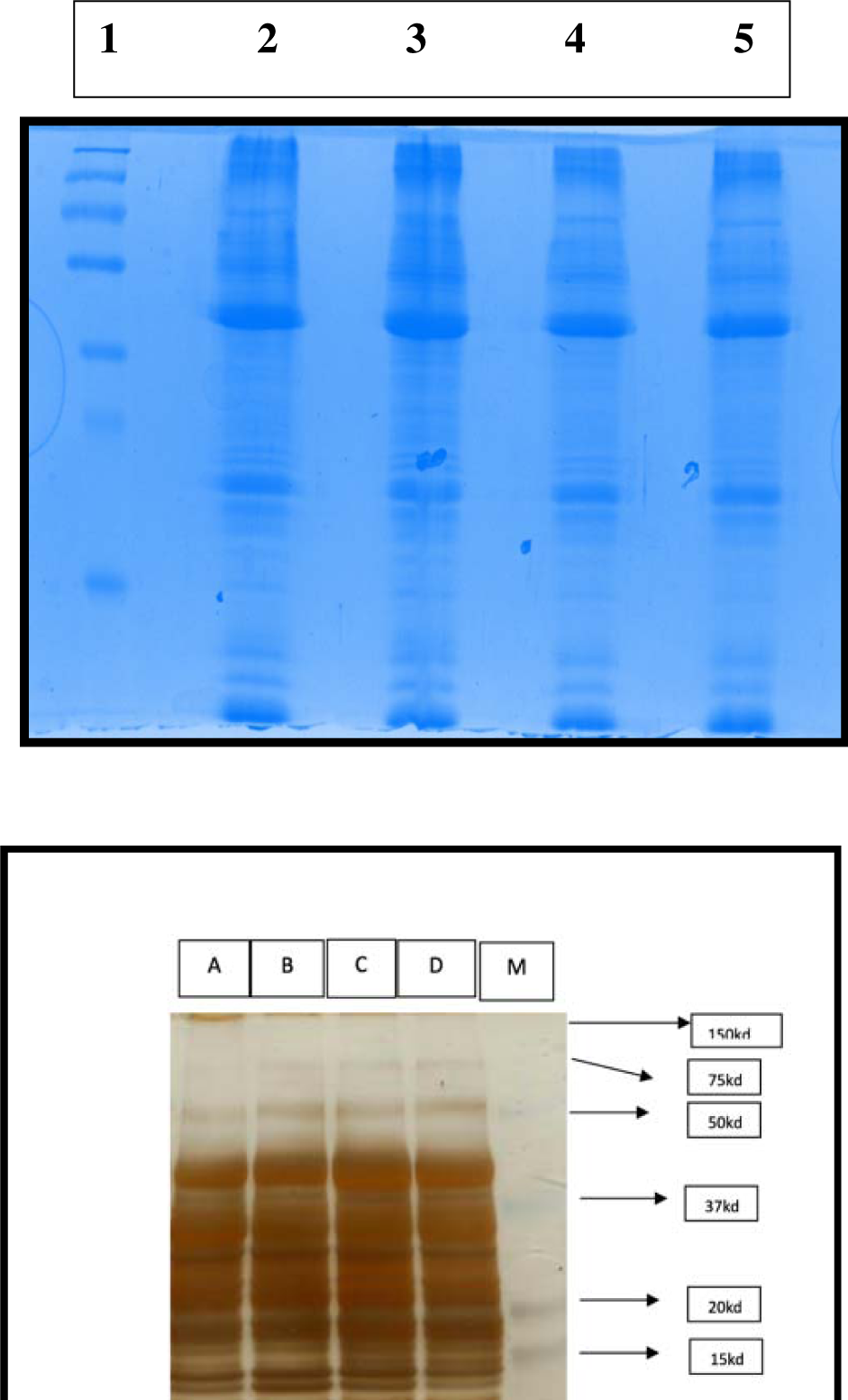

**Supplementary Figure 2.**
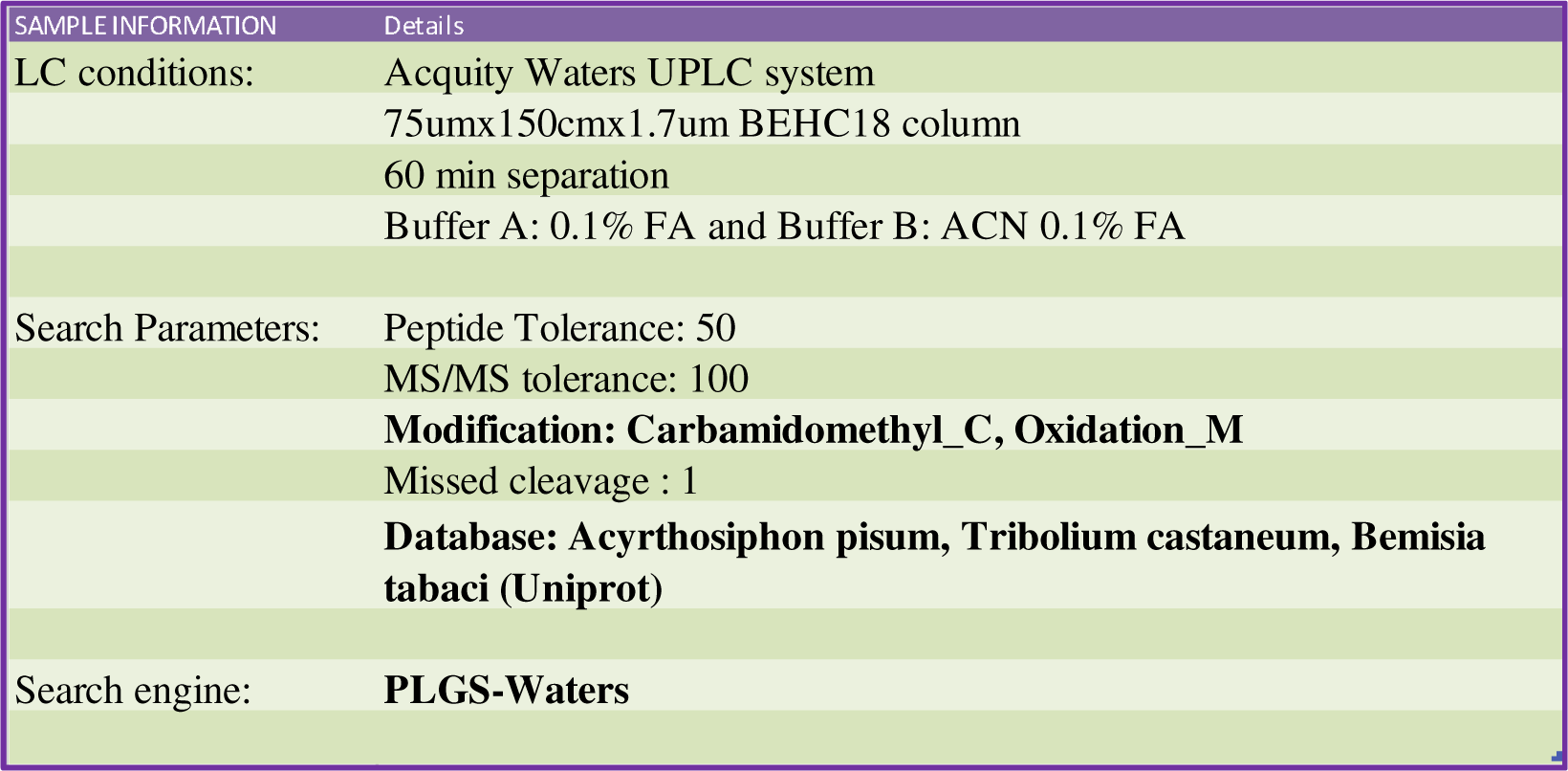
The above table shows the experimental details of LC-MS/MS analysis.

**Supplementary Figure 3.**
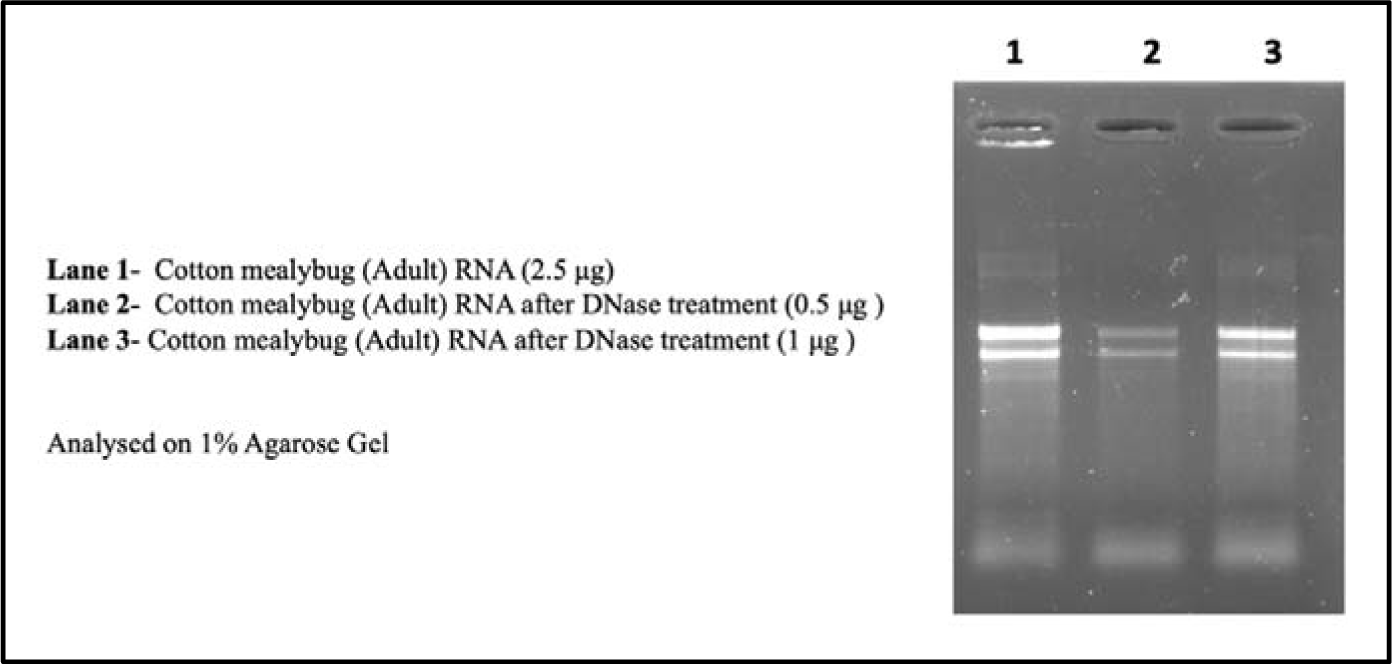
Cotton mealybug (Adult) RNA before and after DNase treatment (on 1% agarose gel).

**Supplementary Figure 4.**
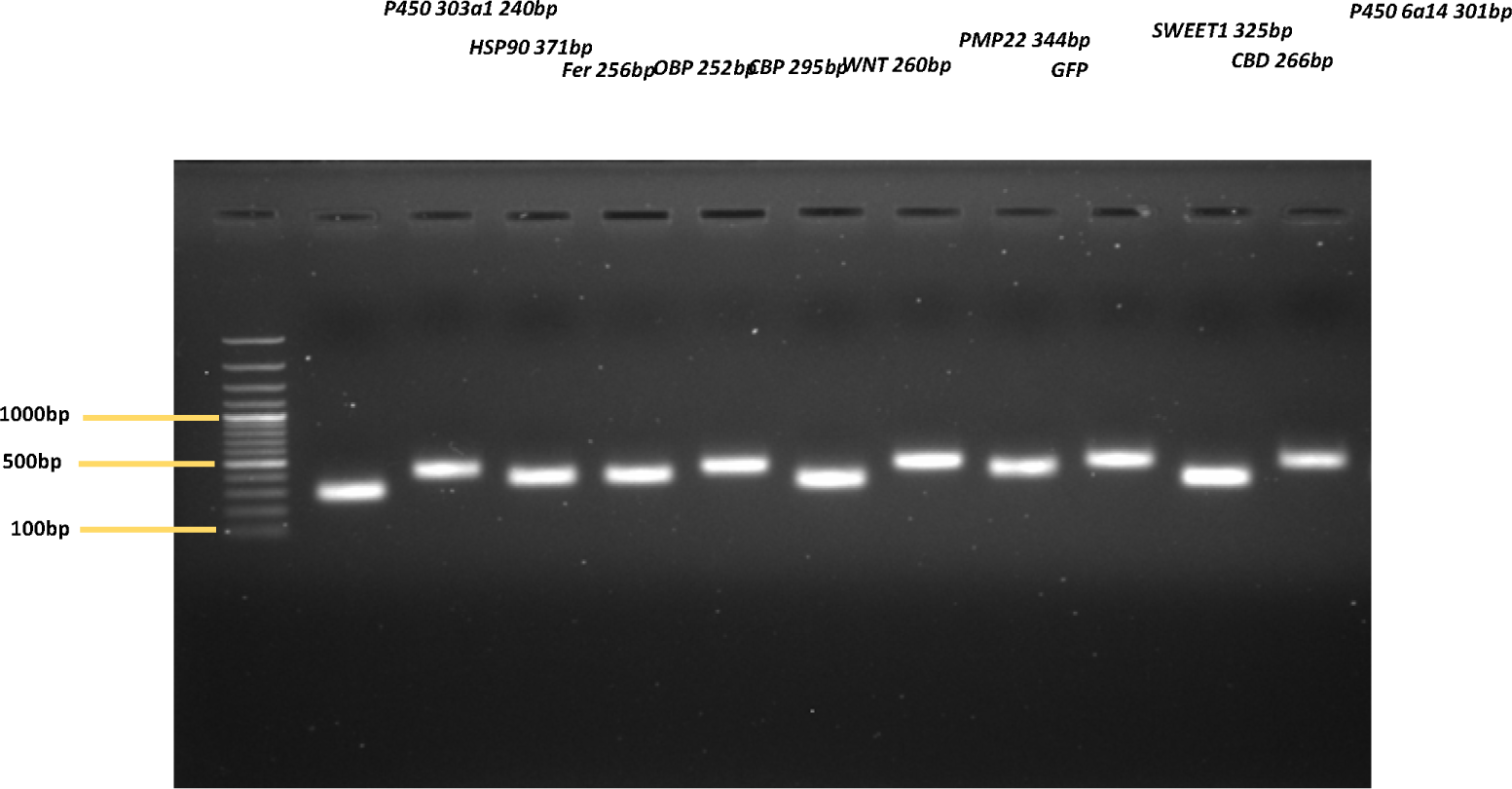
The above gel image is showing PCR amplification of gene targets finalised for oral delivery of dsRNA.

**Supplementary Figure 5.**
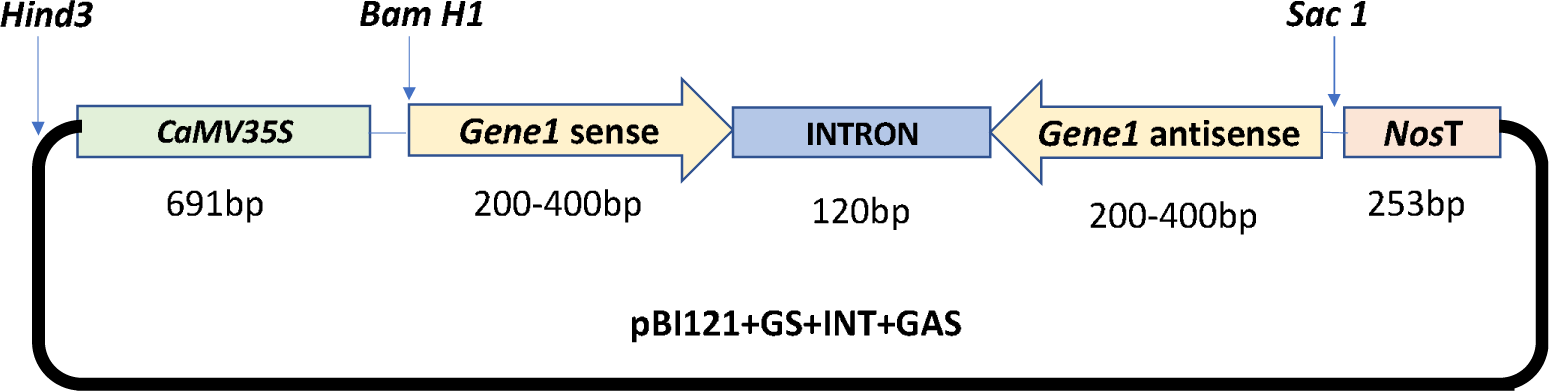
Schematic representation of the gene structure of dsRNA expression cassette. Note: Here, P= CamV35S promoter; S= Sense orientation of the target fragment; I=intron fragments taken from AtRTM1 (120bp); AS= anti-sense orientation of the target fragment.

**Supplementary Table 1.**
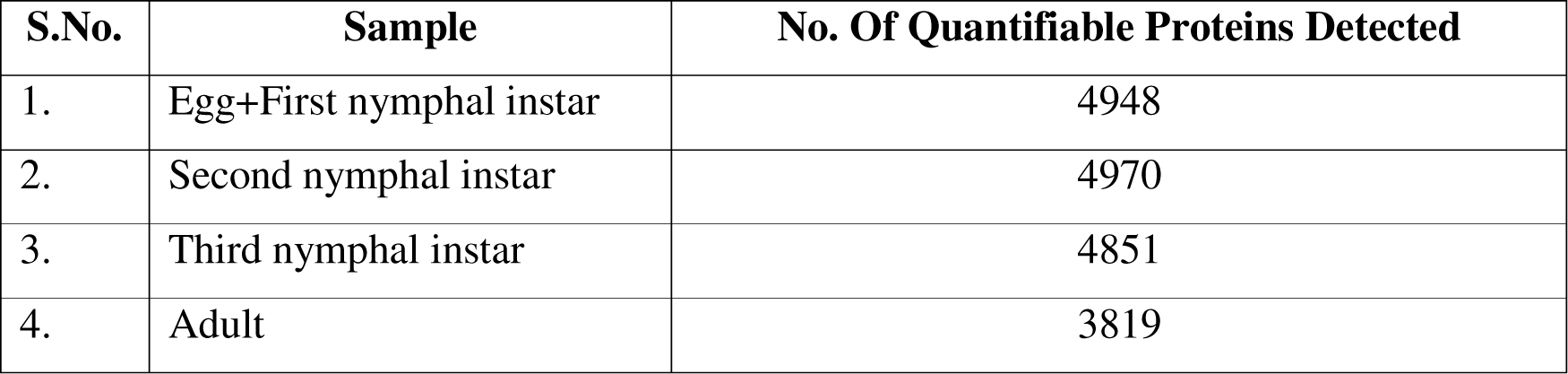
The above table is showing number of DEPs obtained in every group.

**Supplementary Table 2.**
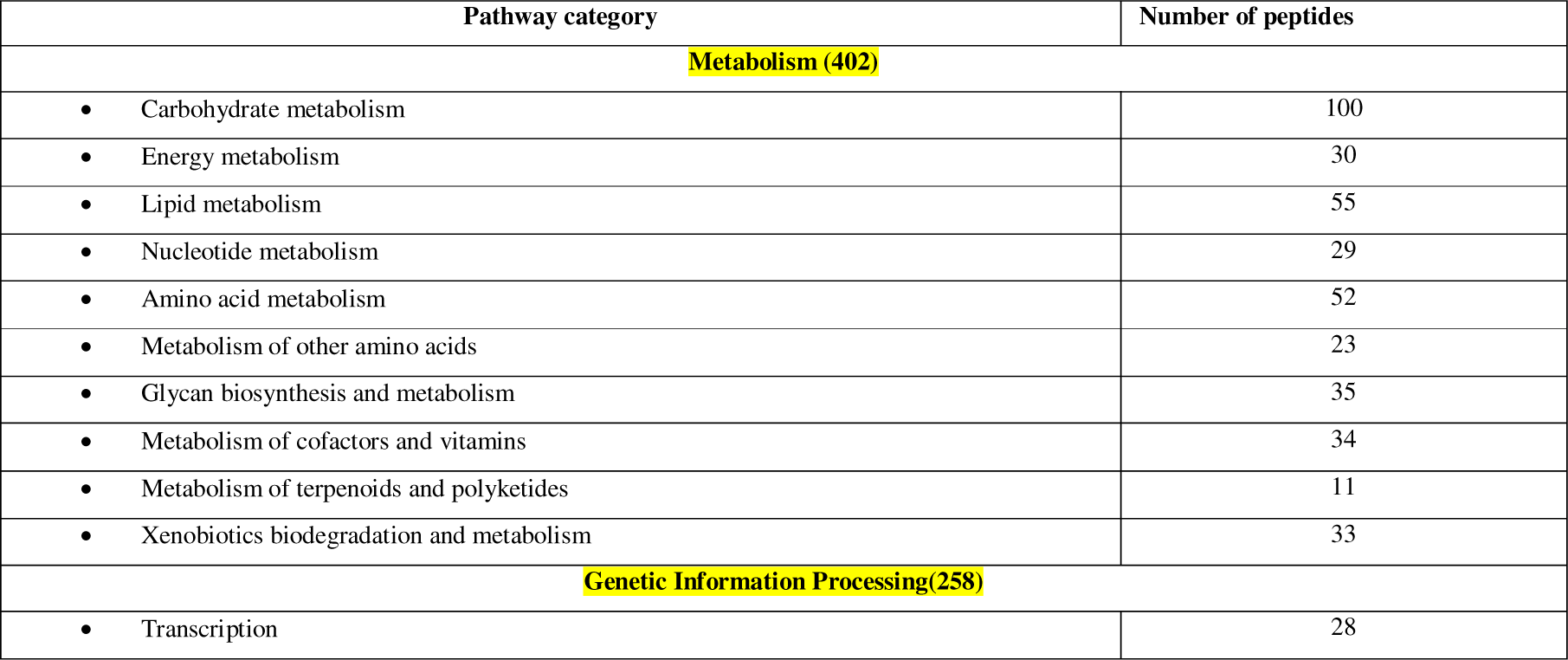
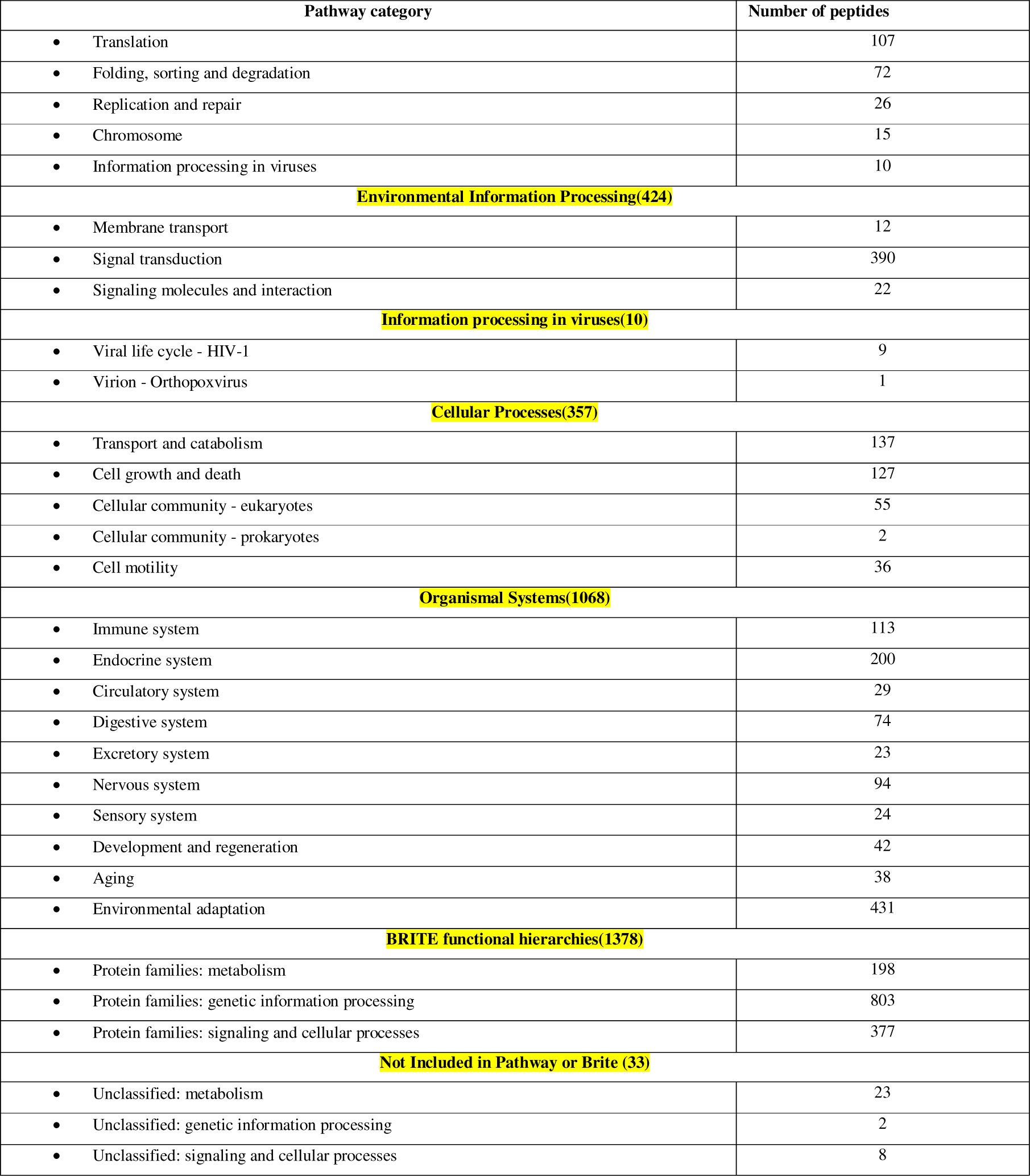
The above table is showing a detailed Kyoto Encyclopedia of Genes and Genomes (KEGG) pathway enrichment analysis of DEPs.

**Supplementary Table 3.**
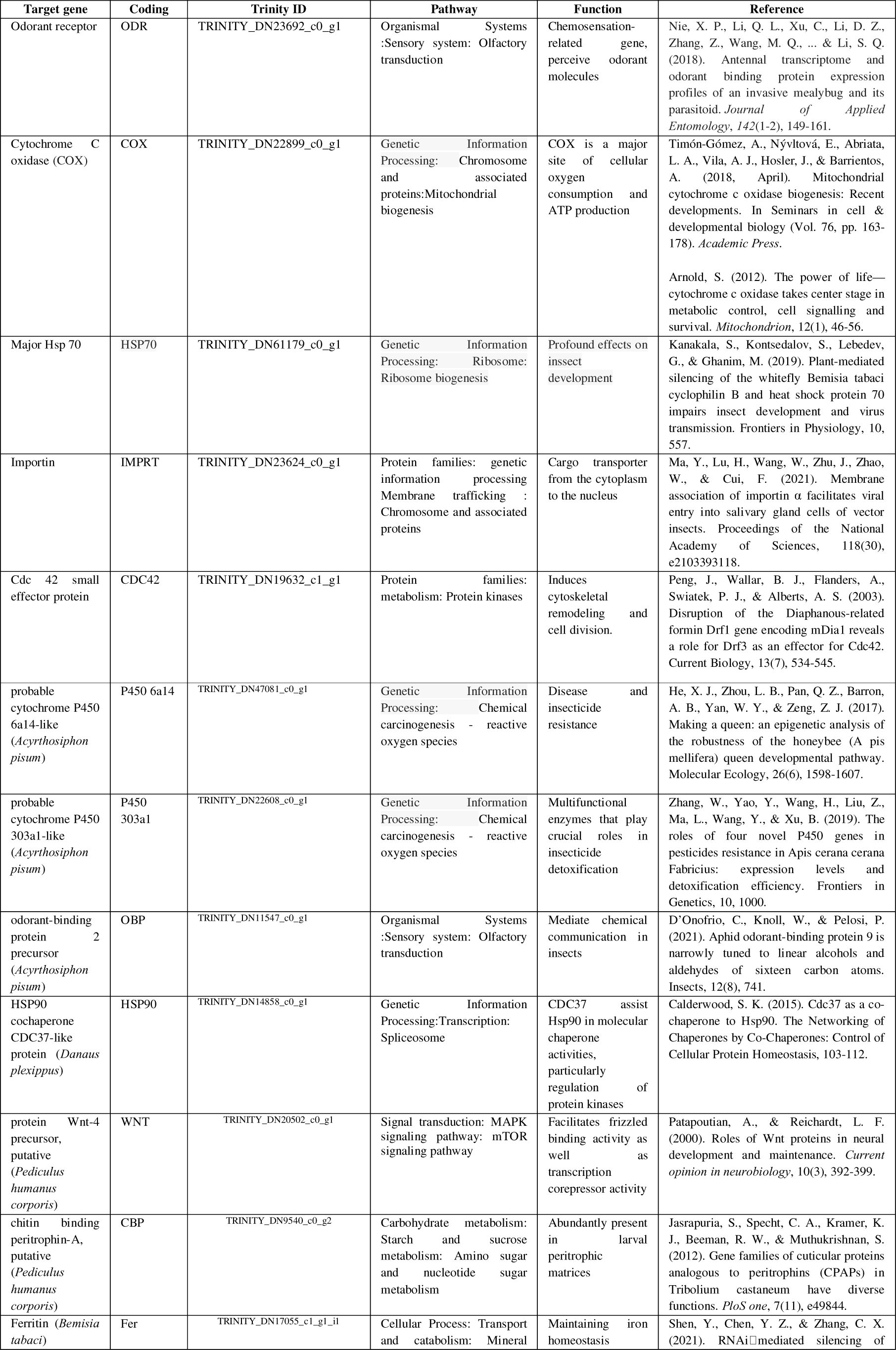
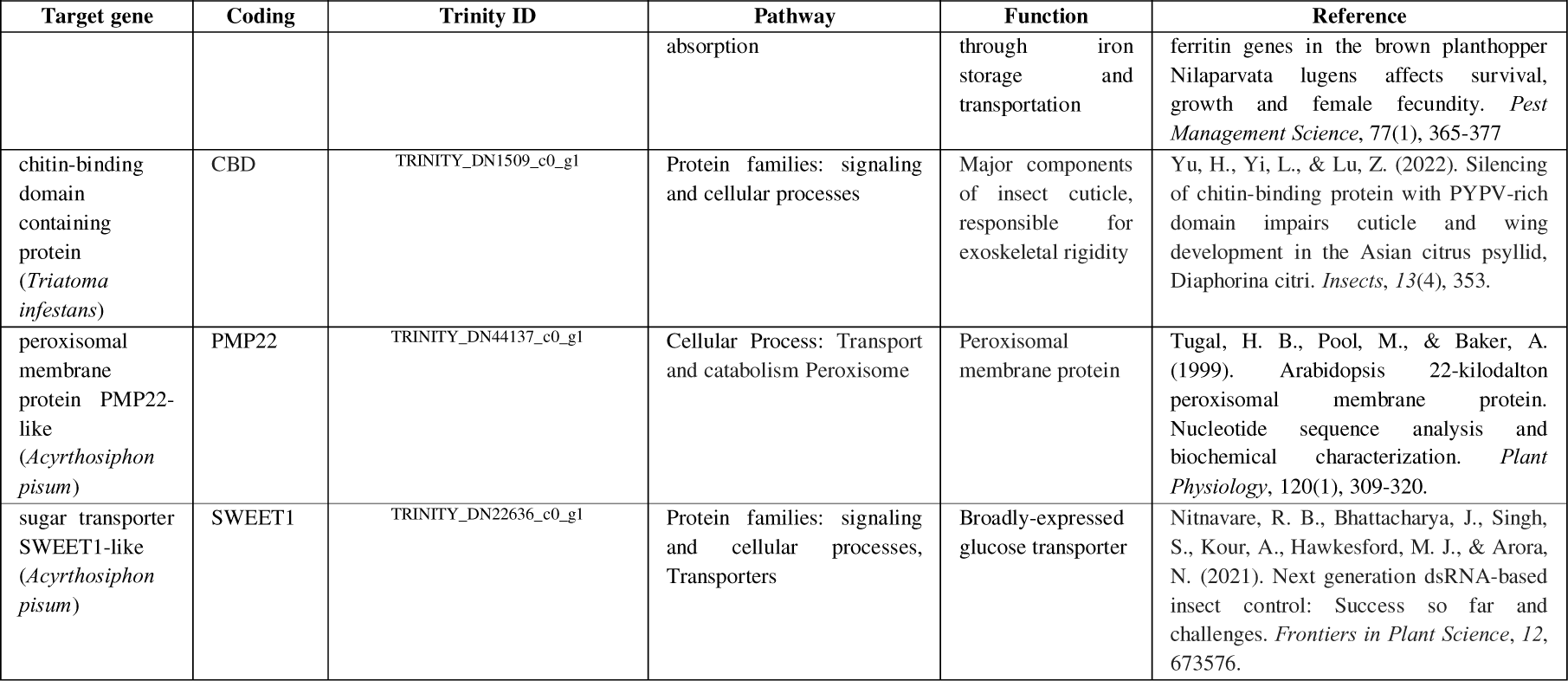
The above table is showing the details of the genes selected for dsRNA feeding bioassays by soaking method.

**Supplementary list 1.**
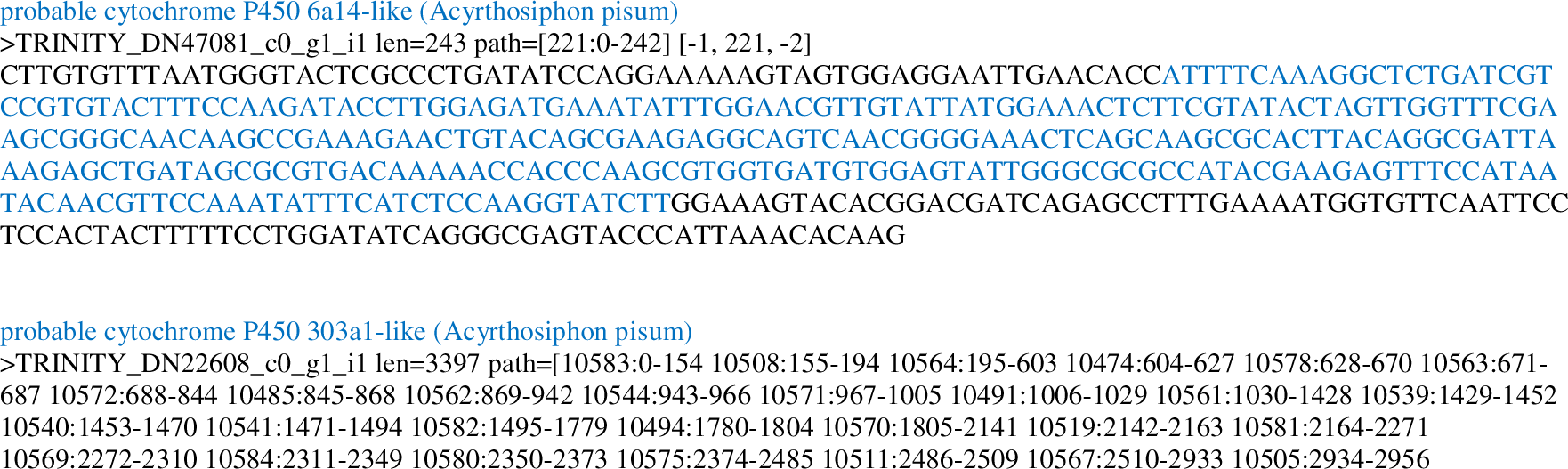
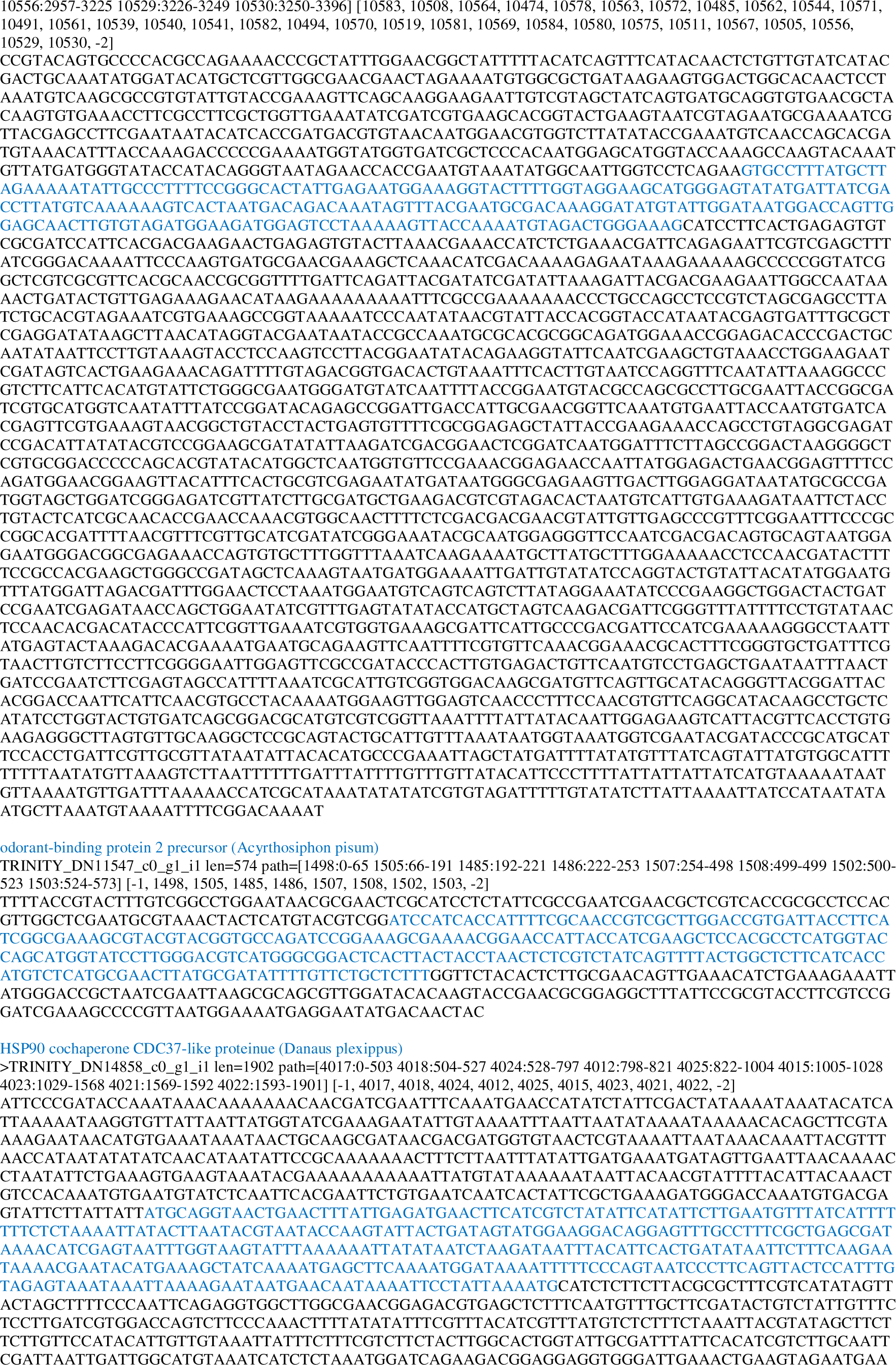
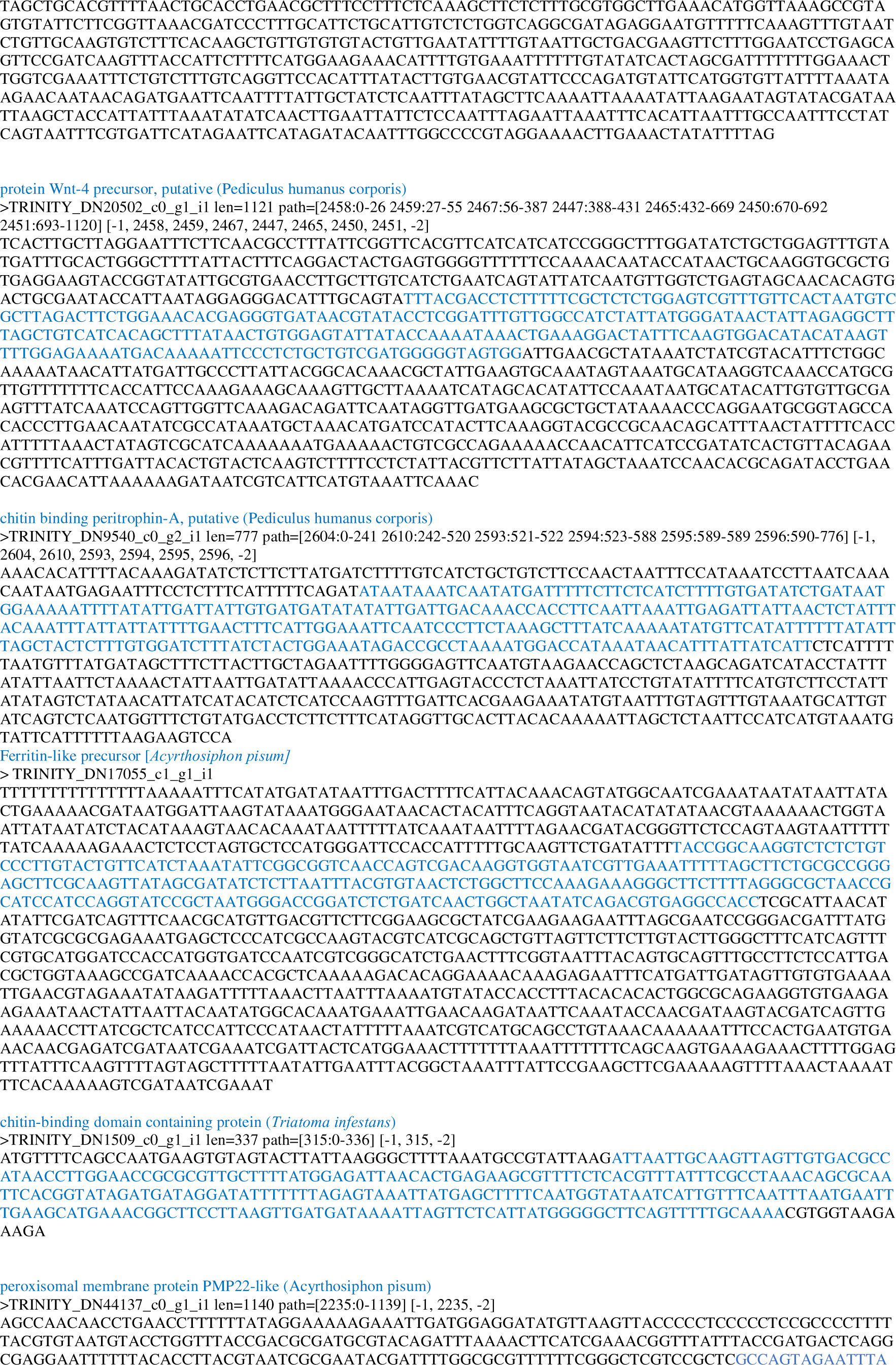
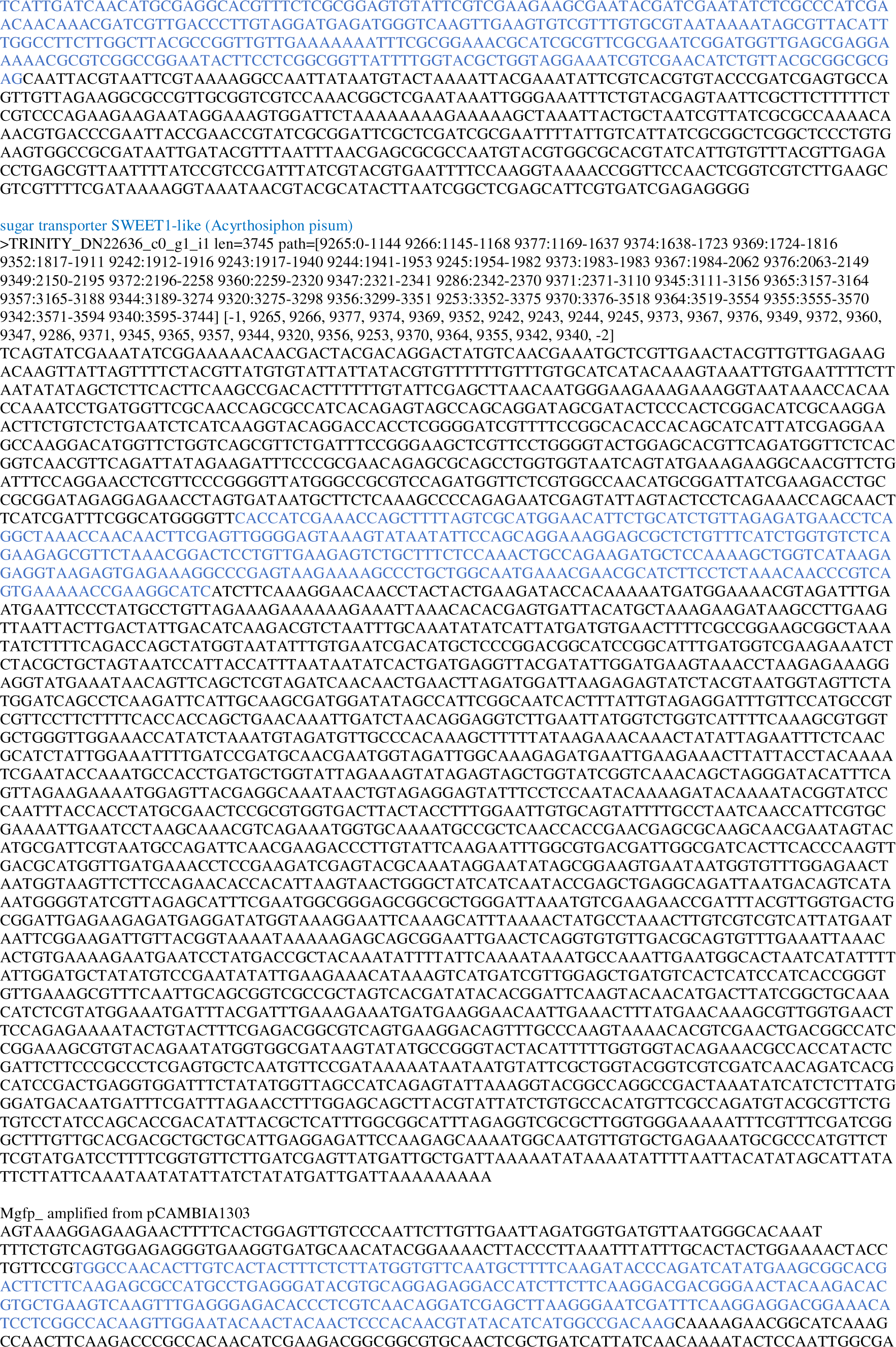
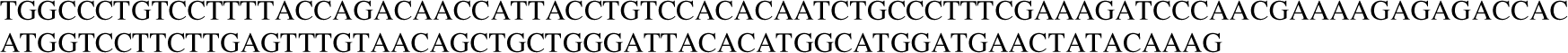
Given below is the list of the genes selected for dsRNA feeding bioassays by oral delivery.

