## SUPPLEMENTAL FILE for "PROTEOME DATA BASED IDENTIFICATION OF POTENTIAL RNAi TARGETS FOR COTTON MEALYBUG (*Phenacoccus solenopsis* Tinsley) POPULATION MANAGEMENT"

**Supplementary Figure 1a :** Pageruler prestained protein ladder, 10 to 180 kDa 3uL of ruler was loaded, **2 -** Adult mealybug protein, (conc.- 10ug/uL) 5uL of sample was loaded with β-ME2%, **3-**Adult mealybug protein, (conc.- 10ug/uL) 5uL of sample was loaded with β-ME 4%**4 -** Adult mealybug protein, (conc.- 10ug/uL) 5uL of sample was loaded with DTT 5mM, **5 -**Adult mealybug protein, (conc.- 10ug/uL) 5uL of sample was loaded with DTT 10mM.

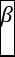

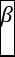

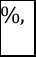

**1	2	3	4	5**

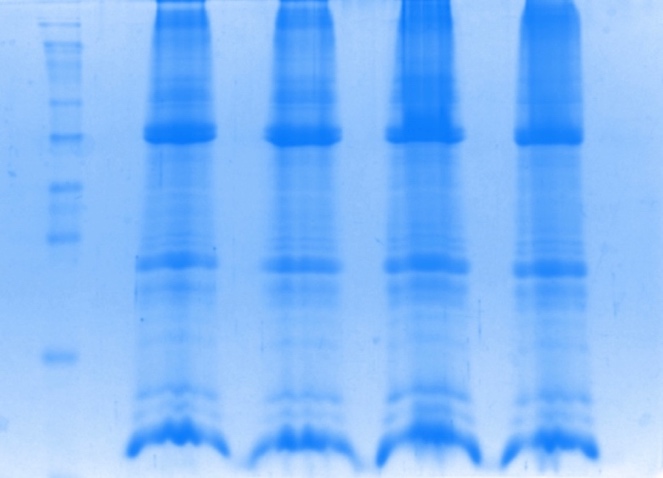

**Supplementary Figure 1b:** Pageruler Prestained Protein Ladder, 10 to 180 kDa 3uL of ruler was loaded, **2 -** Adult mealybug protein, (conc.- 10ug/uL) 3uL of sample was loaded, **3-** 3^rd^ instar mealybug protein, (conc.- 10ug/uL) 3uL of sample**4 –** 2^nd^ instar mealybug protein, (conc.- 10ug/uL) 3uL of sample was loaded, **5-** 1^st^ + Egg mealybug protein, (conc.- 10ug/uL) 3uL of sample.

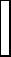

**1	2	3	4	5**

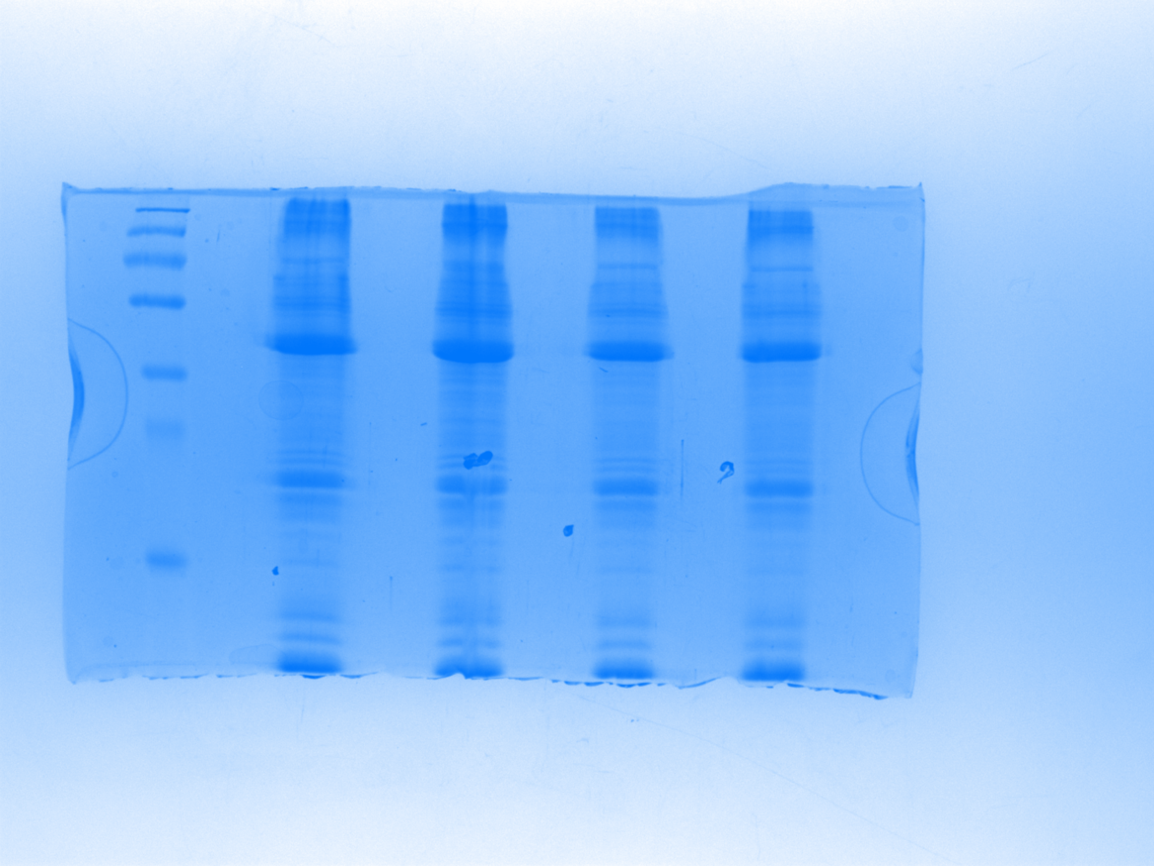

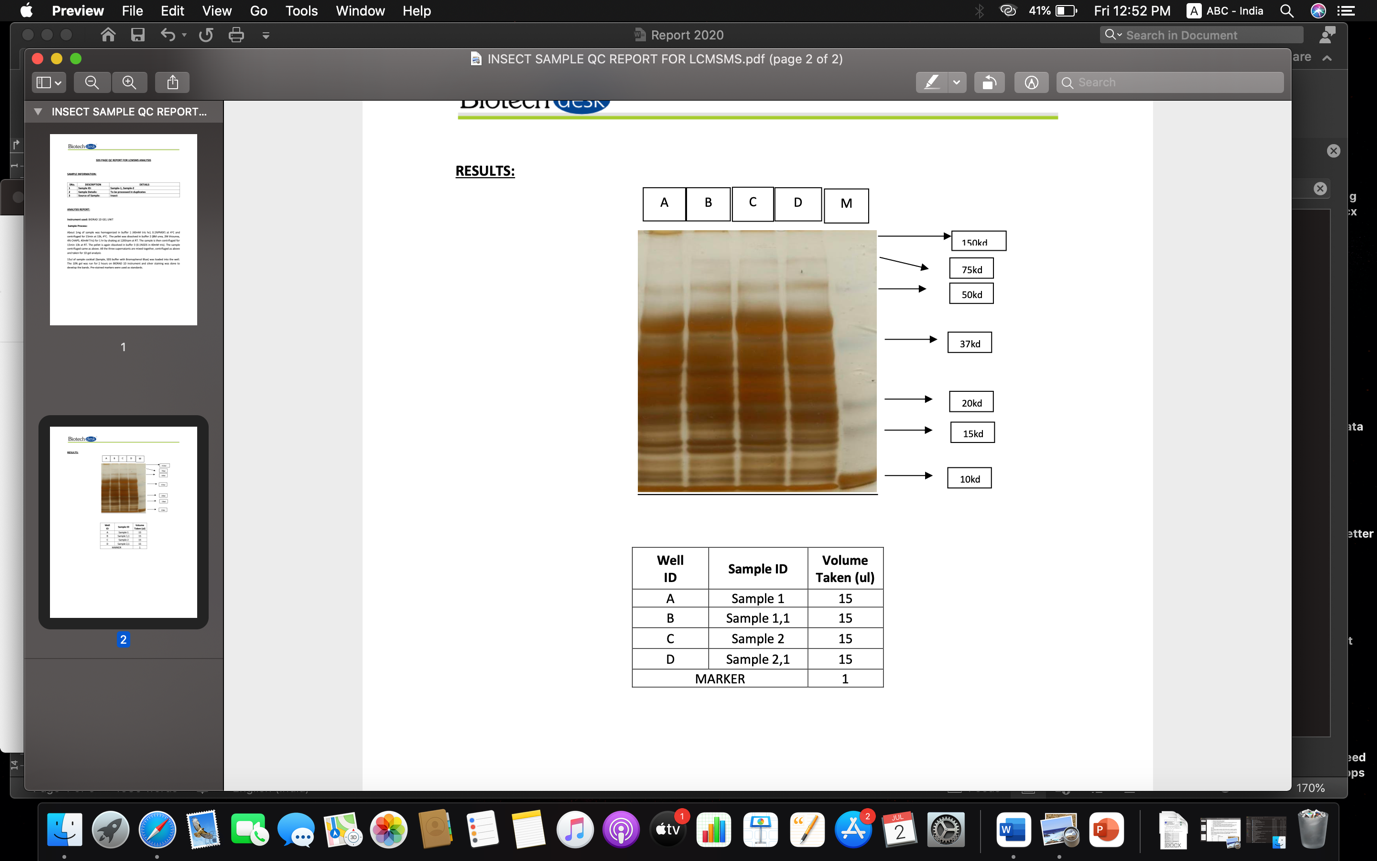

**Supplementary Figure 1c:** Equal amounts of insect protein (60 μg) were analysed on SDS-PAGE (10%) Lane A has protein of adult stage, lane B has 3^rd^ instar, lane C has 2^nd^ instar, and lane D has 1^st^ instart + egg of *P. solenopsis* and lane M has pre-stained marker.

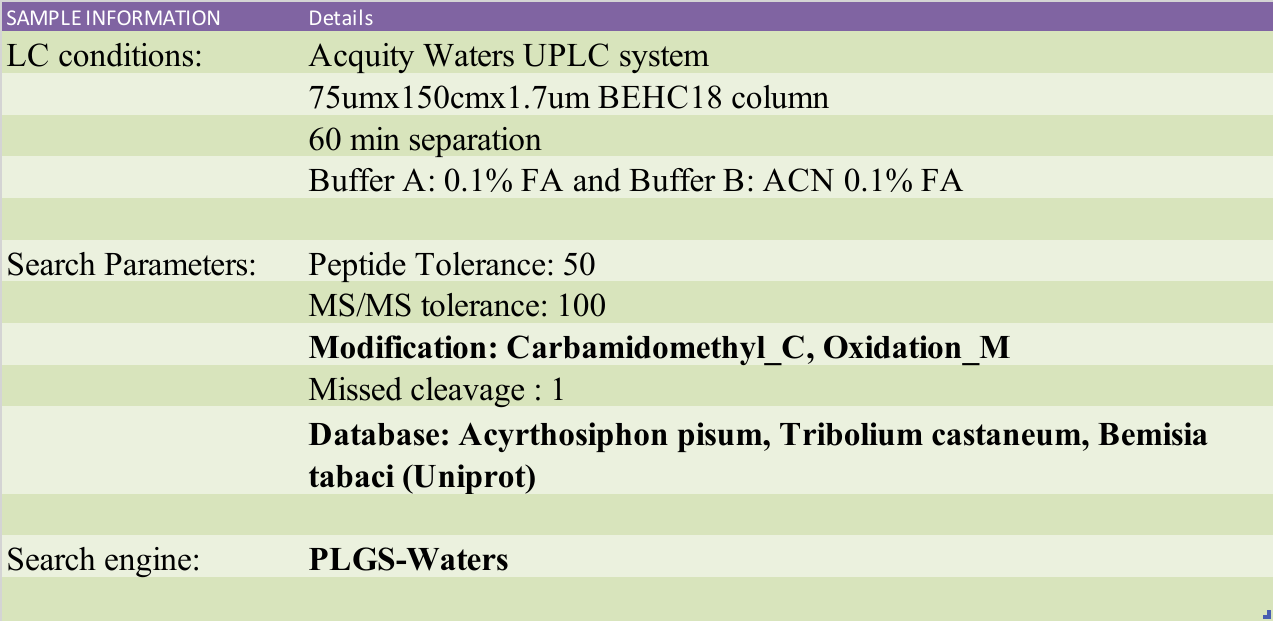

**Supplementary Figure 2. The above table shows the experimental details of LC-MS/MS analysis.**

| **S.No.** | **Sample** | **No. Of Quantifiable Proteins Detected** |
| --- | --- | --- |
| 1. | Egg+First nymphal instar | 4948 |
| 2. | Second nymphal instar | 4970 |
| 3. | Third nymphal instar | 4851 |
| 4. | Adult | 3819 |

**Supplementary Table 1. The above table is showing number of DEPs obtained in every group.**

| **Pathway category** | **Number of peptides** |
| --- | --- |
| **Metabolism (402)** | |
| - Carbohydrate metabolism | 100 |
| - Energy metabolism | 30 |
| - Lipid metabolism | 55 |
| - Nucleotide metabolism | 29 |
| - Amino acid metabolism | 52 |
| - Metabolism of other amino acids | 23 |
| - Glycan biosynthesis and metabolism | 35 |
| - Metabolism of cofactors and vitamins | 34 |
| - Metabolism of terpenoids and polyketides | 11 |
| - Xenobiotics biodegradation and metabolism | 33 |
| **Genetic Information Processing(258)** | |
| - Transcription | 28 |
| - Translation | 107 |
| - Folding, sorting and degradation | 72 |
| - Replication and repair | 26 |
| - Chromosome | 15 |
| - Information processing in viruses | 10 |
| **Environmental Information Processing(424)** | |
| - Membrane transport | 12 |
| - Signal transduction | 390 |
| - Signaling molecules and interaction | 22 |
| **Information processing in viruses(10)** | |
| - Viral life cycle - HIV-1 | 9 |
| - Virion - Orthopoxvirus | 1 |
| **Cellular Processes(357)** | |
| - Transport and catabolism | 137 |
| - Cell growth and death | 127 |
| - Cellular community - eukaryotes | 55 |
| - Cellular community - prokaryotes | 2 |
| - Cell motility | 36 |
| **Organismal Systems(1068)** | |
| - Immune system | 113 |
| - Endocrine system | 200 |
| - Circulatory system | 29 |
| - Digestive system | 74 |
| - Excretory system | 23 |
| - Nervous system | 94 |
| - Sensory system | 24 |
| - Development and regeneration | 42 |
| - Aging | 38 |
| - Environmental adaptation | 431 |
| **BRITE functional hierarchies(1378)** | |
| - Protein families: metabolism | 198 |
| - Protein families: genetic information processing | 803 |
| - Protein families: signaling and cellular processes | 377 |
| **Not Included in Pathway or Brite (33)** | |
| - Unclassified: metabolism | 23 |
| - Unclassified: genetic information processing | 2 |
| - Unclassified: signaling and cellular processes | 8 |

**Supplementary Table 2. The above table is showing a detailed Kyoto Encyclopedia of Genes and Genomes (KEGG) pathway enrichment analysis of DEPs.**

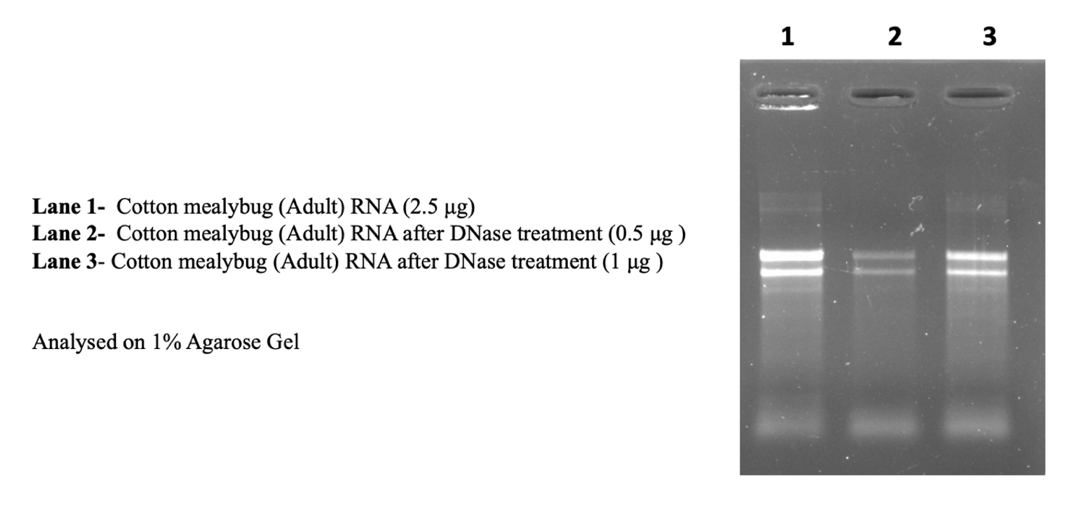

**Supplementary Figure 3. Cotton mealybug (Adult) RNA before and after DNase treatment (on 1% agarose gel).**

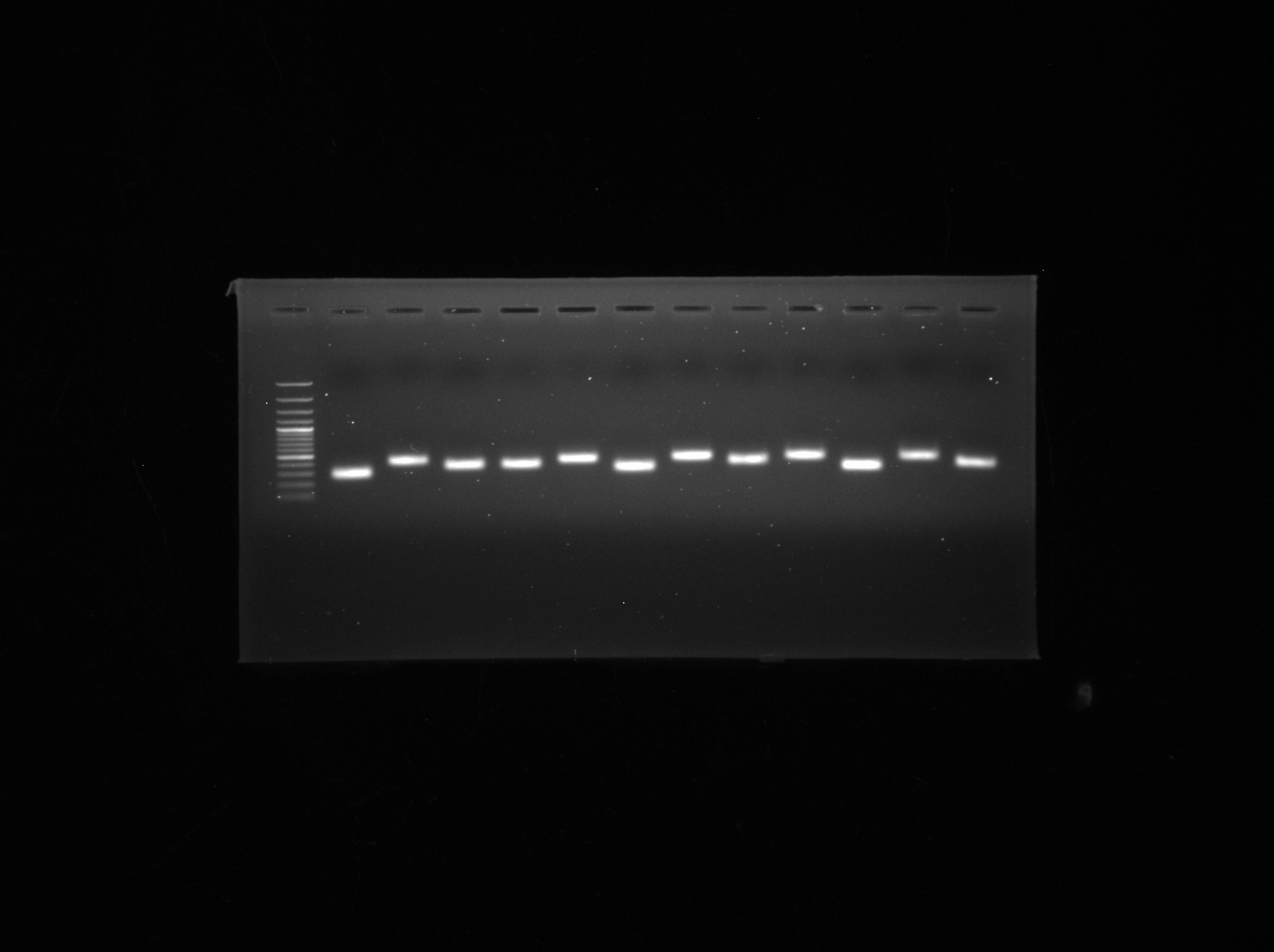

***P450 6a14 301bp***

***OBP 252bp***

***P450 303a1 240bp***

***WNT 260bp***

***Fer 256bp***

***CBP 295bp***

***HSP90 371bp***

***PMP22 344bp***

***GFP 300bp***

***SWEET1 325bp***

***CBD 266bp***

**100bp**

**500bp**

**1000bp**

**Supplementary Figure 4. The above gel image is showing PCR amplification of gene targets finalised for oral delivery of dsRNA.**

| **Target gene** | **Coding** | **Trinity ID** | **Pathway** | **Function** | **Reference** |
| --- | --- | --- | --- | --- | --- |
| Odorant receptor | ODR | TRINITY_DN23692_c0_g1 | Organismal Systems :Sensory system: Olfactory transduction | Chemosensation-related gene, perceive odorant molecules | Nie, X. P., Li, Q. L., Xu, C., Li, D. Z., Zhang, Z., Wang, M. Q., ... & Li, S. Q. (2018). Antennal transcriptome and odorant binding protein expression profiles of an invasive mealybug and its parasitoid. *Journal of Applied Entomology*, *142*(1-2), 149-161. |
| Cytochrome C oxidase (COX) | COX | TRINITY_DN22899_c0_g1 | Genetic Information Processing: Chromosome and associated proteins:Mitochondrial biogenesis | COX is a major site of cellular oxygen consumption and ATP production | Timón-Gómez, A., Nývltová, E., Abriata, L. A., Vila, A. J., Hosler, J., & Barrientos, A. (2018, April). Mitochondrial cytochrome c oxidase biogenesis: Recent developments. In Seminars in cell & developmental biology (Vol. 76, pp. 163-178). *Academic* *Press*.  Arnold, S. (2012). The power of life—cytochrome c oxidase takes center stage in metabolic control, cell signalling and survival. *Mitochondrion*, 12(1), 46-56. |
| Major Hsp 70 | HSP70 | TRINITY_DN61179_c0_g1 | Genetic Information Processing: Ribosome: Ribosome biogenesis | Profound effects on inssect development | Kanakala, S., Kontsedalov, S., Lebedev, G., & Ghanim, M. (2019). Plant-mediated silencing of the whitefly Bemisia tabaci cyclophilin B and heat shock protein 70 impairs insect development and virus transmission. Frontiers in Physiology, 10, 557. |
| Importin | IMPRT | TRINITY_DN23624_c0_g1 | Protein families: genetic information processing Membrane trafficking : Chromosome and associated proteins | Cargo transporter from the cytoplasm to the nucleus | Ma, Y., Lu, H., Wang, W., Zhu, J., Zhao, W., & Cui, F. (2021). Membrane association of importin α facilitates viral entry into salivary gland cells of vector insects. Proceedings of the National Academy of Sciences, 118(30), e2103393118. |
| Cdc 42 small effector protein | CDC42 | TRINITY_DN19632_c1_g1 | Protein families: metabolism: Protein kinases | Induces cytoskeletal remodeling and cell division. | Peng, J., Wallar, B. J., Flanders, A., Swiatek, P. J., & Alberts, A. S. (2003). Disruption of the Diaphanous-related formin Drf1 gene encoding mDia1 reveals a role for Drf3 as an effector for Cdc42. Current Biology, 13(7), 534-545. |
| probable cytochrome P450 6a14-like (*Acyrthosiphon pisum*) | P450 6a14 | TRINITY_DN47081_c0_g1 | Genetic Information Processing: Chemical carcinogenesis - reactive oxygen species | Disease and insecticide resistance | He, X. J., Zhou, L. B., Pan, Q. Z., Barron, A. B., Yan, W. Y., & Zeng, Z. J. (2017). Making a queen: an epigenetic analysis of the robustness of the honeybee (A pis mellifera) queen developmental pathway. Molecular Ecology, 26(6), 1598-1607. |
| probable cytochrome P450 303a1-like (*Acyrthosiphon pisum*) | P450 303a1 | TRINITY_DN22608_c0_g1 | Genetic Information Processing: Chemical carcinogenesis - reactive oxygen species | Multifunctional enzymes that play crucial roles in insecticide detoxification | Zhang, W., Yao, Y., Wang, H., Liu, Z., Ma, L., Wang, Y., & Xu, B. (2019). The roles of four novel P450 genes in pesticides resistance in Apis cerana cerana Fabricius: expression levels and detoxification efficiency. Frontiers in Genetics, 10, 1000. |
| odorant-binding protein 2 precursor (*Acyrthosiphon pisum*) | OBP | TRINITY_DN11547_c0_g1 | Organismal Systems :Sensory system: Olfactory transduction | Mediate chemical communication in insects | D’Onofrio, C., Knoll, W., & Pelosi, P. (2021). Aphid odorant-binding protein 9 is narrowly tuned to linear alcohols and aldehydes of sixteen carbon atoms. Insects, 12(8), 741. |
| HSP90 cochaperone CDC37-like protein (*Danaus plexippus*) | HSP90 | TRINITY_DN14858_c0_g1 | Genetic Information Processing:Transcription: Spliceosome | CDC37 assist Hsp90 in molecular chaperone activities, particularly regulation of protein kinases | Calderwood, S. K. (2015). Cdc37 as a co-chaperone to Hsp90. The Networking of Chaperones by Co-Chaperones: Control of Cellular Protein Homeostasis, 103-112. |
| protein Wnt-4 precursor, putative (*Pediculus humanus corporis*) | WNT | TRINITY_DN20502_c0_g1 | Signal transduction: MAPK signaling pathway: mTOR signaling pathway | Facilitates frizzled binding activity as well as transcription corepressor activity | Patapoutian, A., & Reichardt, L. F. (2000). Roles of Wnt proteins in neural development and maintenance. *Current opinion in neurobiology*, 10(3), 392-399. |
| chitin binding peritrophin-A, putative (*Pediculus humanus corporis*) | CBP | TRINITY_DN9540_c0_g2 | Carbohydrate metabolism: Starch and sucrose metabolism: Amino sugar and nucleotide sugar metabolism | Abundantly present in larval peritrophic matrices | Jasrapuria, S., Specht, C. A., Kramer, K. J., Beeman, R. W., & Muthukrishnan, S. (2012). Gene families of cuticular proteins analogous to peritrophins (CPAPs) in Tribolium castaneum have diverse functions. *PloS one*, 7(11), e49844. |
| Ferritin (*Bemisia tabaci*) | Fer | TRINITY_DN17055_c1_g1_i1 | Cellular Process: Transport and catabolism: Mineral absorption | Maintaining iron homeostasis through iron storage and transportation | Shen, Y., Chen, Y. Z., & Zhang, C. X. (2021). RNAi‐mediated silencing of ferritin genes in the brown planthopper Nilaparvata lugens affects survival, growth and female fecundity. *Pest Management Science*, 77(1), 365-377 |
| chitin-binding domain containing protein (*Triatoma infestans*) | CBD | TRINITY_DN1509_c0_g1 | Protein families: signaling and cellular processes | Major components of insect cuticle, responsible for exoskeletal rigidity | Yu, H., Yi, L., & Lu, Z. (2022). Silencing of chitin-binding protein with PYPV-rich domain impairs cuticle and wing development in the Asian citrus psyllid, Diaphorina citri. *Insects*, *13*(4), 353. |
| peroxisomal membrane protein PMP22-like (*Acyrthosiphon pisum*) | PMP22 | TRINITY_DN44137_c0_g1 | Cellular Process: Transport and catabolism Peroxisome | Peroxisomal membrane protein | Tugal, H. B., Pool, M., & Baker, A. (1999). Arabidopsis 22-kilodalton peroxisomal membrane protein. Nucleotide sequence analysis and biochemical characterization. *Plant Physiology*, 120(1), 309-320. |
| sugar transporter SWEET1-like (*Acyrthosiphon pisum*) | SWEET1 | TRINITY_DN22636_c0_g1 | Protein families: signaling and cellular processes, Transporters | Broadly-expressed glucose transporter | Nitnavare, R. B., Bhattacharya, J., Singh, S., Kour, A., Hawkesford, M. J., & Arora, N. (2021). Next generation dsRNA-based insect control: Success so far and challenges. *Frontiers in Plant Science*, *12*, 673576. |

**Supplementary Table 3. The above table is showing the details of the genes selected for dsRNA feeding bioassays by soaking method.**

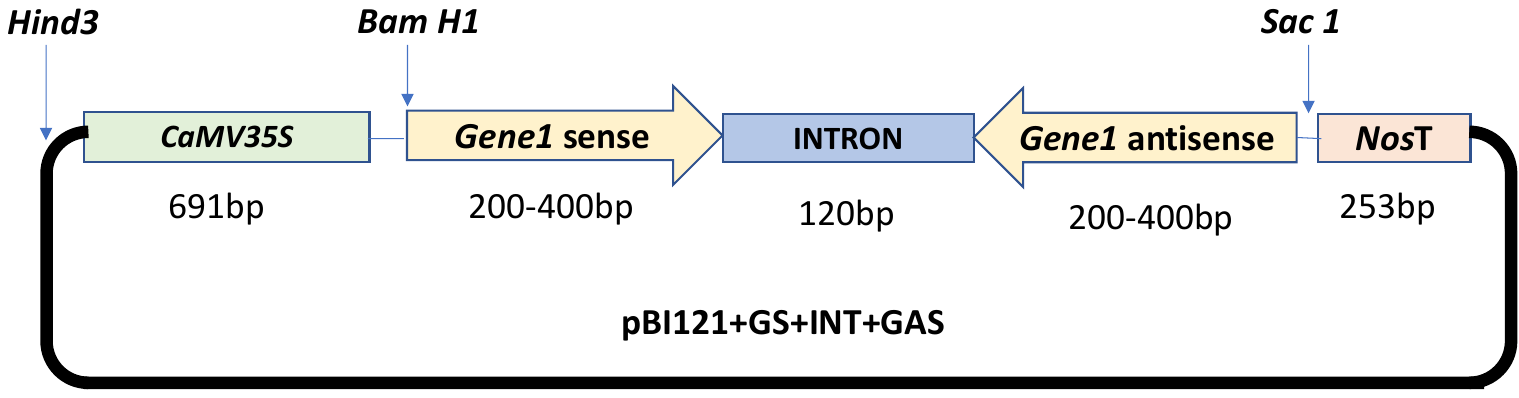

**Supplementary Figure 5. Schematic representation of the gene structure of dsRNA expression cassette.**

Note: Here, P= CamV35S promoter; S= Sense orientation of the target fragment; I=intron fragments taken from AtRTM1 (120bp); AS= anti-sense orientation of the target fragment.

**Supplementary list 1. Given below is the list of the genes selected for dsRNA feeding bioassays by oral delivery.**

probable cytochrome P450 6a14-like (Acyrthosiphon pisum)

>TRINITY_DN47081_c0_g1_i1 len=243 path=[221:0-242] [-1, 221, -2]

CTTGTGTTTAATGGGTACTCGCCCTGATATCCAGGAAAAAGTAGTGGAGGAATTGAACACCATTTTCAAAGGCTCTGATCGTCCGTGTACTTTCCAAGATACCTTGGAGATGAAATATTTGGAACGTTGTATTATGGAAACTCTTCGTATACTAGTTGGTTTCGAAGCGGGCAACAAGCCGAAAGAACTGTACAGCGAAGAGGCAGTCAACGGGGAAACTCAGCAAGCGCACTTACAGGCGATTAAAGAGCTGATAGCGCGTGACAAAAACCACCCAAGCGTGGTGATGTGGAGTATTGGGCGCGCCATACGAAGAGTTTCCATAATACAACGTTCCAAATATTTCATCTCCAAGGTATCTTGGAAAGTACACGGACGATCAGAGCCTTTGAAAATGGTGTTCAATTCCTCCACTACTTTTTCCTGGATATCAGGGCGAGTACCCATTAAACACAAG

probable cytochrome P450 303a1-like (Acyrthosiphon pisum)

>TRINITY_DN22608_c0_g1_i1 len=3397 path=[10583:0-154 10508:155-194 10564:195-603 10474:604-627 10578:628-670 10563:671-687 10572:688-844 10485:845-868 10562:869-942 10544:943-966 10571:967-1005 10491:1006-1029 10561:1030-1428 10539:1429-1452 10540:1453-1470 10541:1471-1494 10582:1495-1779 10494:1780-1804 10570:1805-2141 10519:2142-2163 10581:2164-2271 10569:2272-2310 10584:2311-2349 10580:2350-2373 10575:2374-2485 10511:2486-2509 10567:2510-2933 10505:2934-2956 10556:2957-3225 10529:3226-3249 10530:3250-3396] [10583, 10508, 10564, 10474, 10578, 10563, 10572, 10485, 10562, 10544, 10571, 10491, 10561, 10539, 10540, 10541, 10582, 10494, 10570, 10519, 10581, 10569, 10584, 10580, 10575, 10511, 10567, 10505, 10556, 10529, 10530, -2]

CCGTACAGTGCCCCACGCCAGAAAACCCGCTATTTGGAACGGCTATTTTTACATCAGTTTCATACAACTCTGTTGTATCATACGACTGCAAATATGGATACATGCTCGTTGGCGAACGAACTAGAAAATGTGGCGCTGATAAGAAGTGGACTGGCACAACTCCTAAATGTCAAGCGCCGTGTATTGTACCGAAAGTTCAGCAAGGAAGAATTGTCGTAGCTATCAGTGATGCAGGTGTGAACGCTACAAGTGTGAAACCTTCGCCTTCGCTGGTTGAAATATCGATCGTGAAGCACGGTACTGAAGTAATCGTAGAATGCGAAAATCGTTACGAGCCTTCGAATAATACATCACCGATGACGTGTAACAATGGAACGTGGTCTTATATACCGAAATGTCAACCAGCACGATGTAAACATTTACCAAAGACCCCCGAAAATGGTATGGTGATCGCTCCCACAATGGAGCATGGTACCAAAGCCAAGTACAAATGTTATGATGGGTATACCATACAGGGTAATAGAACCACCGAATGTAAATATGGCAATTGGTCCTCAGAAGTGCCTTTATGCTTAGAAAAATATTGCCCTTTTCCGGGCACTATTGAGAATGGAAAGGTACTTTTGGTAGGAAGCATGGGAGTATATGATTATCGACCTTATGTCAAAAAAGTCACTAATGACAGACAAATAGTTTACGAATGCGACAAAGGATATGTATTGGATAATGGACCAGTTGGAGCAACTTGTGTAGATGGAAGATGGAGTCCTAAAAAGTTACCAAAATGTAGACTGGGAAAGCATCCTTCACTGAGAGTGTCGCGATCCATTCACGACGAAGAACTGAGAGTGTACTTAAACGAAACCATCTCTGAAACGATTCAGAGAATTCGTCGAGCTTTATCGGGACAAAATTCCCAAGTGATGCGAACGAAAGCTCAAACATCGACAAAAGAGAATAAAGAAAAAGCCCCCGGTATCGGCTCGTCGCGTTCACGCAACCGCGGTTTTGATTCAGATTACGATATCGATATTAAAGATTACGACGAAGAATTGGCCAATAAAACTGATACTGTTGAGAAAGAACATAAGAAAAAAAAATTTCGCCGAAAAAAACCCTGCCAGCCTCCGTCTAGCGAGCCTTATCTGCACGTAGAAATCGTGAAAGCCGGTAAAAATCCCAATATAACGTATTACCACGGTACCATAATACGAGTGATTTGCGCTCGAGGATATAAGCTTAACATAGGTACGAATAATACCGCCAAATGCGCACGCGGCAGATGGAAACCGGAGACACCCGACTGCAATATAATTCCTTGTAAAGTACCTCCAAGTCCTTACGGAATATACAGAAGGTATTCAATCGAAGCTGTAAACCTGGAAGAATCGATAGTCACTGAAGAAACAGATTTTGTAGACGGTGACACTGTAAATTTCACTTGTAATCCAGGTTTCAATATTAAAGGCCCGTCTTCATTCACATGTATTCTGGGCGAATGGGATGTATCAATTTTACCGGAATGTACGCCAGCGCCTTGCGAATTACCGGCGATCGTGCATGGTCAATATTTATCCGGATACAGAGCCGGATTGACCATTGCGAACGGTTCAAATGTGAATTACCAATGTGATCACGAGTTCGTGAAAGTAACGGCTGTACCTACTGAGTGTTTTCGCGGAGAGCTATTACCGAAGAAACCAGCCTGTAGGCGAGATCCGACATTATATACGTCCGGAAGCGATATATTAAGATCGACGGAACTCGGATCAATGGATTTCTTAGCCGGACTAAGGGGCTCGTGCGGACCCCCAGCACGTATACATGGCTCAATGGTGTTCCGAAACGGAGAACCAATTATGGAGACTGAACGGAGTTTTCCAGATGGAACGGAAGTTACATTTCACTGCGTCGAGAATATGATAATGGGCGAGAAGTTGACTTGGAGGATAATATGCGCCGATGGTAGCTGGATCGGGAGATCGTTATCTTGCGATGCTGAAGACGTCGTAGACACTAATGTCATTGTGAAAGATAATTCTACCTGTACTCATCGCAACACCGAACCAAACGTGGCAACTTTTCTCGACGACGAACGTATTGTTGAGCCCGTTTCGGAATTTCCCGCCGGCACGATTTTAACGTTTCGTTGCATCGATATCGGGAAATACGCAATGGAGGGTTCCAATCGACGACAGTGCAGTAATGGAGAATGGGACGGCGAGAAACCAGTGTGCTTTGGTTTAAATCAAGAAAATGCTTATGCTTTGGAAAAACCTCCAACGATACTTTTCCGCCACGAAGCTGGGCCGATAGCTCAAAGTAATGATGGAAAATTGATTGTATATCCAGGTACTGTATTACATATGGAATGTTTATGGATTAGACGATTTGGAACTCCTAAATGGAATGTCAGTCAGTCTTATAGGAAATATCCCGAAGGCTGGACTACTGATCCGAATCGAGATAACCAGCTGGAATATCGTTTGAGTATATACCATGCTAGTCAAGACGATTCGGGTTTATTTTCCTGTATAACTCCAACACGACATACCCATTCGGTTGAAATCGTGGTGAAAGCGATTCATTGCCCGACGATTCCATCGAAAAAGGGCCTAATTATGAGTACTAAAGACACGAAAATGAATGCAGAAGTTCAATTTTCGTGTTCAAACGGAAACGCACTTTCGGGTGCTGATTTCGTAACTTGTCTTCCTTCGGGGAATTGGAGTTCGCCGATACCCACTTGTGAGACTGTTCAATGTCCTGAGCTGAATAATTTAACTGATCCGAATCTTCGAGTAGCCATTTTAAATCGCATTGTCGGTGGACAAGCGATGTTCAGTTGCATACAGGGTTACGGATTACACGGACCAATTCATTCAACGTGCCTACAAAATGGAAGTTGGAGTCAACCCTTTCCAACGTGTTCAGGCATACAAGCCTGCTCATATCCTGGTACTGTGATCAGCGGACGCATGTCGTCGGTTAAATTTTATTATACAATTGGAGAAGTCATTACGTTCACCTGTGAAGAGGGCTTAGTGTTGCAAGGCTCCGCAGTACTGCATTGTTTAAATAATGGTAAATGGTCGAATACGATACCCGCATGCATTCCACCTGATTCGTTGCGTTATAATATTACACATGCCCGAAATTAGCTATGATTTTATATGTTTATCAGTATTATGTGGCATTTTTTTTAATATGTTAAAGTCTTAATTTTTTGATTTATTTTGTTTGTTATACATTCCCTTTTATTATTATTATCATGTAAAAATAATGTTAAAATGTTGATTTAAAAACCATCGCATAAATATATATCGTGTAGATTTTTGTATATCTTATTAAAATTATCCATAATATAATGCTTAAATGTAAAATTTTCGGACAAAAT

odorant-binding protein 2 precursor (Acyrthosiphon pisum)

TRINITY_DN11547_c0_g1_i1 len=574 path=[1498:0-65 1505:66-191 1485:192-221 1486:222-253 1507:254-498 1508:499-499 1502:500-523 1503:524-573] [-1, 1498, 1505, 1485, 1486, 1507, 1508, 1502, 1503, -2]

TTTTACCGTACTTTGTCGGCCTGGAATAACGCGAACTCGCATCCTCTATTCGCCGAATCGAACGCTCGTCACCGCGCCTCCACGTTGGCTCGAATGCGTAAACTACTCATGTACGTCGGATCCATCACCATTTTCGCAACCGTCGCTTGGACCGTGATTACCTTCATCGGCGAAAGCGTACGTACGGTGCCAGATCCGGAAAGCGAAAACGGAACCATTACCATCGAAGCTCCACGCCTCATGGTACCAGCATGGTATCCTTGGGACGTCATGGGCGGACTCACTTACTACCTAACTCTCGTCTATCAGTTTTACTGGCTCTTCATCACCATGTCTCATGCGAACTTATGCGATATTTTGTTCTGCTCTTTGGTTCTACACTCTTGCGAACAGTTGAAACATCTGAAAGAAATTATGGGACCGCTAATCGAATTAAGCGCAGCGTTGGATACACAAGTACCGAACGCGGAGGCTTTATTCCGCGTACCTTCGTCCGGATCGAAAGCCCCGTTAATGGAAAATGAGGAATATGACAACTAC

HSP90 cochaperone CDC37-like proteinue (Danaus plexippus)

>TRINITY_DN14858_c0_g1_i1 len=1902 path=[4017:0-503 4018:504-527 4024:528-797 4012:798-821 4025:822-1004 4015:1005-1028 4023:1029-1568 4021:1569-1592 4022:1593-1901] [-1, 4017, 4018, 4024, 4012, 4025, 4015, 4023, 4021, 4022, -2]

ATTCCCGATACCAAATAAACAAAAAAACAACGATCGAATTTCAAATGAACCATATCTATTCGACTATAAAATAAATACATCATTAAAAATAAGGTGTTATTAATTATGGTATCGAAAGAATATTGTAAAATTTAATTAATATAAAATAAAAACACAGCTTCGTAAAAGAATAACATGTGAAATAAATAACTGCAAGCGATAACGACGATGGTGTAACTCGTAAAATTAATAAACAAATTACGTTTAACCATAATATATATCAACATAATATTCCGCAAAAAAACTTTCTTAATTTATATTGATGAAATGATAGTTGAATTAACAAAACCTAATATTCTGAAAGTGAAGTAAATACGAAAAAAAAAAATTATGTATAAAAAATAATTACAACGTATTTTACATTACAAACTGTCCACAAATGTGAATGTATCTCAATTCACGAATTCTGTGAATCAATCACTATTCGCTGAAAGATGGGACCAAATGTGACGAGTATTCTTATTATTATGCAGGTAACTGAACTTTATTGAGATGAACTTCATCGTCTATATTCATATTCTTGAATGTTTATCATTTTTTTCTCTAAAATTATACTTAATACGTAATACCAAGTATTACTGATAGTATGGAAGGACAGGAGTTTGCCTTTCGCTGAGCGATAAAACATCGAGTAATTTGGTAAGTATTTAAAAAATTATATAATCTAAGATAATTTACATTCACTGATATAATTCTTTCAAGAATAAAACGAATACATGAAAGCTATCAAAATGAGCTTCAAAATGGATAAAATTTTTCCCAGTAATCCCTTCAGTTACTCCATTTGTAGAGTAAATAAATTAAAAGAATAATGAACAATAAAATTCCTATTAAAATGCATCTCTTCTTACGCGCTTTCGTCATATAGTTACTAGCTTTTCCCAATTCAGAGGTGGCTTGGCGAACGGAGACGTGAGCTCTTTCAATGTTTGCTTCGATACTGTCTATTGTTTCTCCTTGATCGTGGACCAGTCTTCCCAAACTTTTATATATTTCGTTTACATCGTTTATGTCTCTTTCTAAATTACGTATAGCTTCTTCTTGTTCCATACATTGTTGTAAATTATTTCTTTCGTCTTCTACTTGGCACTGGTATTGCGATTTATTCACATCGTCTTGCAATTCGATTAATTGATTGGCATGTAAATCATCTCTAAATGGATCAGAAGACGGAGGAGGTGGGATTGAAACTGAAGTAGAATGAATAGCTGCACGTTTTAACTGCACCTGAACGCTTTCCTTTCTCAAAGCTTCTCTTTGCGTGGCTTGAAACATGGTTAAAGCCGTAGTGTATTCTTCGGTTAAACGATCCCTTTGCATTCTGCATTGTCTCTGGTCAGGCGATAGAGGAATGTTTTTCAAAGTTTGTAATCTGTTGCAAGTGTCTTTCACAAGCTGTTGTGTGTACTGTTGAATATTTTGTAATTGCTGACGAAGTTCTTTGGAATCCTGAGCAGTTCCGATCAAGTTTACCATTCTTTTCATGGAAGAAACATTTTGTGAAATTTTTTGTATATCACTAGCGATTTTTTTGGAAACTTGGTCGAAATTTCTGTCTTTGTCAGGTTCCACATTTATACTTGTGAACGTATTCCCAGATGTATTCATGGTGTTATTTTAAATAAGAACAATAACAGATGAATTCAATTTTATTGCTATCTCAATTTATAGCTTCAAAATTAAAATATTAAGAATAGTATACGATAATTAAGCTACCATTATTTAAATATATCAACTTGAATTATTCTCCAATTTAGAATTAAATTTCACATTAATTTGCCAATTTCCTATCAGTAATTTCGTGATTCATAGAATTCATAGATACAATTTGGCCCCGTAGGAAAACTTGAAACTATATTTTAG

protein Wnt-4 precursor, putative (Pediculus humanus corporis)

>TRINITY_DN20502_c0_g1_i1 len=1121 path=[2458:0-26 2459:27-55 2467:56-387 2447:388-431 2465:432-669 2450:670-692 2451:693-1120] [-1, 2458, 2459, 2467, 2447, 2465, 2450, 2451, -2]

TCACTTGCTTAGGAATTTCTTCAACGCCTTTATTCGGTTCACGTTCATCATCATCCGGGCTTTGGATATCTGCTGGAGTTTGTATGATTTGCACTGGGCTTTTATTACTTTCAGGACTACTGAGTGGGGTTTTTTCCAAAACAATACCATAACTGCAAGGTGCGCTGTGAGGAAGTACCGGTATATTGCGTGAACCTTGCTTGTCATCTGAATCAGTATTATCAATGTTGGTCTGAGTAGCAACACAGTGACTGCGAATACCATTAATAGGAGGGACATTTGCAGTATTTACGACCTCTTTTTCGCTCTCTGGAGTCGTTTGTTCACTAATGTCGCTTAGACTTCTGGAAACACGAGGGTGATAACGTATACCTCGGATTTGTTGGCCATCTATTATGGGATAACTATTAGAGGCTTTAGCTGTCATCACAGCTTTATAACTGTGGAGTATTATACCAAAATAAACTGAAAGGACTATTTCAAGTGGACATACATAAGTTTTGGAGAAAATGACAAAAATTCCCTCTGCTGTCGATGGGGGTAGTGGATTGAACGCTATAAATCTATCGTACATTTCTGGCAAAAATAACATTATGATTGCCCTTATTACGGCACAAACGCTATTGAAGTGCAAATAGTAAATGCATAAGGTCAAACCATGCGTTGTTTTTTTCACCATTCCAAAGAAAGCAAAGTTGCTTAAAATCATAGCACATATTCCAAATAATGCATACATTGTGTTGCGAAGTTTATCAAATCCAGTTGGTTCAAAGACAGATTCAATAGGTTGATGAAGCGCTGCTATAAAACCCAGGAATGCGGTAGCCACACCCTTGAACAATATCGCCATAAATGCTAAACATGATCCATACTTCAAAGGTACGCCGCAACAGCATTTAACTATTTTCACCATTTTTAAACTATAGTCGCATCAAAAAAATGAAAAACTGTCGCCAGAAAAACCAACATTCATCCGATATCACTGTTACAGAACGTTTTCATTTGATTACACTGTACTCAAGTCTTTTCCTCTATTACGTTCTTATTATAGCTAAATCCAACACGCAGATACCTGAACACGAACATTAAAAAAGATAATCGTCATTCATGTAAATTCAAAC

chitin binding peritrophin-A, putative (Pediculus humanus corporis)

>TRINITY_DN9540_c0_g2_i1 len=777 path=[2604:0-241 2610:242-520 2593:521-522 2594:523-588 2595:589-589 2596:590-776] [-1, 2604, 2610, 2593, 2594, 2595, 2596, -2]

AAACACATTTTACAAAGATATCTCTTCTTATGATCTTTTGTCATCTGCTGTCTTCCAACTAATTTCCATAAATCCTTAATCAAACAATAATGAGAATTTCCTCTTTCATTTTTCAGATATAATAAATCAATATGATTTTTCTTCTCATCTTTTGTGATATCTGATAATGGAAAAATTTTATATTGATTATTGTGATGATATATATTGATTGACAAACCACCTTCAATTAAATTGAGATTATTAACTCTATTTACAAATTTATTATTATTTTGAACTTTCATTGGAAATTCAATCCCTTCTAAAGCTTTATCAAAAATATGTTCATATTTTTTATATTTAGCTACTCTTTGTGGATCTTTATCTACTGGAAATAGACCGCCTAAAATGGACCATAAATAACATTTATTATCATTCTCATTTTTAATGTTTATGATAGCTTTCTTACTTGCTAGAATTTTGGGGAGTTCAATGTAAGAACCAGCTCTAAGCAGATCATACCTATTTATATTAATTCTAAAACTATTAATTGATATTAAAACCCATTGAGTACCCTCTAAATTATCCTGTATATTTTCATGTCTTCCTATTATATAGTCTATAACATTATCATACATCTCATCCAAGTTTGATTCACGAAGAAATATGTAATTTGTAGTTTGTAAATGCATTGTATCAGTCTCAATGGTTTCTGTATGACCTCTTCTTTCATAGGTTGCACTTACACAAAAATTAGCTCTAATTCCATCATGTAAATGTATTCATTTTTTAAGAAGTCCA

Ferritin-like precursor [*Acyrthosiphon pisum]*

> TRINITY_DN17055_c1_g1_i1

TTTTTTTTTTTTTTTAAAAATTTCATATGATATAATTTGACTTTTCATTACAAACAGTATGGCAATCGAAATAATATAATTATACTGAAAAACGATAATGGATTAAGTATAAATGGGAATAACACTACATTTCAGGTAATACATATATAACGTAAAAAACTGGTAATTATAATATCTACATAAAGTAACACAAATAATTTTTATCAAATAATTTTAGAACGATACGGGTTCTCCAGTAAGTAATTTTTTATCAAAAAGAAACTCTCCTAGTGCTCCATGGGATTCCACCATTTTTGCAAGTTCTGATATTTTACCGGCAAGGTCTCTCTGTCCCTTGTACTGTTCATCTAAATATTCGGCGGTCAACCAGTCGACAAGGTGGTAATCGTTGAAATTTTTAGCTTCTGCGCCGGGAGCTTCGCAAGTTATAGCGATATCTCTTAATTTACGTGTAACTCTGGCTTCCAAAGAAAGGGCTTCTTTTAGGGCGCTAACCGCATCCATCCAGGTATCCGCTAATGGGACCGGATCTCTGATCAACTGGCTAATATCAGACGTGAGGCCACCTCGCATTAACATATATTCGATCAGTTTCAACGCATGTTGACGTTCTTCGGAAGCGCTATCGAAGAAGAATTTAGCGAATCCGGGACGATTTATGGTATCGCGCGAGAAATGAGCTCCCATCGCCAAGTACGTCATCGCAGCTGTTAGTTCTTCTTGTACTTGGGCTTTCATCAGTTTCGTGCATGGATCCACCATGGTGATCCAATCGTCGGGCATCTGAACTTTCGGTAATTTACAGTGCAGTTTGCCTTCTCCATTGACGCTGGTAAAGCCGATCAAAACCACGCTCAAAAAGACACAGGAAAACAAAGAGAATTTCATGATTGATAGTTGTGTGAAAATTGAACGTAGAAATATAAGATTTTTAAACTTAATTTAAAATGTATACCACCTTTACACACACTGGCGCAGAAGGTGTGAAGAAGAAATAACTATTAATTACAATATGGCACAAATGAAATTGAACAAGATAATTCAAATACCAACGATAAGTACGATCAGTTGAAAAACCTTATCGCTCATCCATTCCCATAACTATTTTTAAATCGTCATGCAGCCTGTAAACAAAAAATTTCCACTGAATGTGAAACAACGAGATCGATAATCGAAATCGATTACTCATGGAAACTTTTTTTAAATTTTTTTCAGCAAGTGAAAGAAACTTTTGGAGTTTATTTCAAGTTTTAGTAGCTTTTTAATATTGAATTTACGGCTAAATTTATTCCGAAGCTTCGAAAAAGTTTTAAACTAAAATTTCACAAAAAGTCGATAATCGAAAT

chitin-binding domain containing protein (*Triatoma infestans*)

>TRINITY_DN1509_c0_g1_i1 len=337 path=[315:0-336] [-1, 315, -2]

ATGTTTTCAGCCAATGAAGTGTAGTACTTATTAAGGGCTTTTAAATGCCGTATTAAGATTAATTGCAAGTTAGTTGTGACGCCATAACCTTGGAACCGCGCGTTGCTTTTATGGAGATTAACACTGAGAAGCGTTTTCTCACGTTTATTTCGCCTAAACAGCGCAATTCACGGTATAGATGATAGGATATTTTTTTAGAGTAAATTATGAGCTTTTCAATGGTATAATCATTGTTTCAATTTAATGAATTTGAAGCATGAAACGGCTTCCTTAAGTTGATGATAAAATTAGTTCTCATTATGGGGGCTTCAGTTTTTGCAAAACGTGGTAAGAAAGA

peroxisomal membrane protein PMP22-like (Acyrthosiphon pisum)

>TRINITY_DN44137_c0_g1_i1 len=1140 path=[2235:0-1139] [-1, 2235, -2]

AGCCAACAACCTGAACCTTTTTTATAGGAAAAAGAAATTGATGGAGGATATGTTAAGTTACCCCCTCCCCCTCCGCCCCTTTTTACGTGTAATGTACCTGGTTTACCGACGCGATGCGTACAGATTTAAAACTTCATCGAAACGGTTTATTTACCGATGACTCAGGCGAGGAATTTTTTACACCTTACGTAATCGCGAATACGATTTTGGCGCGTTTTTTCGGGCTCGTCCGCTCGCCAGTAGAATTTATCATTGATCAACATGCGAGGCACGTTTCTCGCGGAGTGTATTCGTCGAAGAAGCGAATACGATCGAATATCTCGCCCATCGAACAACAAACGATCGTTGACCCTTGTAGGATGAGATGGGTCAAGTTGAAGTGTCGTTTGTGCGTAATAAAATAGCGTTACATTTGGCCTTCTTGGCTTACGCCGGTTGTTGAAAAAAATTTCGCGGAAACGCATCGCGTTCGCGAATCGGATGGTTGAGCGAGGAAAAACGCGTCGGCCGGAATACTTCCTCGGCGGTTATTTTGGTACGCTGGTAGGAAATCGTCGAACATCTGTTACGCGGCGCGAGCAATTACGTAATTCGTAAAAGGCCAATTATAATGTACTAAAATTACGAAATATTCGTCACGTGTACCCGATCGAGTGCCAGTTGTTAGAAGGCGCCGTTGCGGTCGTCCAAACGGCTCGAATAAATTGGGAAATTTCTGTACGAGTAATTCGCTTCTTTTTCTCGTCCCAGAAGAAGAATAGGAAAGTGGATTCTAAAAAAAAGAAAAAGCTAAATTACTGCTAATCGTTATCGCGCCAAAACAAACGTGACCCGAATTACCGAACCGTATCGCGGATTCGCTCGATCGCGAATTTTATTGTCATTATCGCGGCTCGGCTCCCTGTGAAGTGGCCGCGATAATTGATACGTTTAATTTAACGAGCGCGCCAATGTACGTGGCGCACGTATCATTGTGTTTACGTTGAGACCTGAGCGTTAATTTTATCCGTCCGATTTATCGTACGTGAATTTTCCAAGGTAAAACCGGTTCCAACTCGGTCGTCTTGAAGCGTCGTTTTCGATAAAAGGTAAATAACGTACGCATACTTAATCGGCTCGAGCATTCGTGATCGAGAGGGG

sugar transporter SWEET1-like (Acyrthosiphon pisum)

>TRINITY_DN22636_c0_g1_i1 len=3745 path=[9265:0-1144 9266:1145-1168 9377:1169-1637 9374:1638-1723 9369:1724-1816 9352:1817-1911 9242:1912-1916 9243:1917-1940 9244:1941-1953 9245:1954-1982 9373:1983-1983 9367:1984-2062 9376:2063-2149 9349:2150-2195 9372:2196-2258 9360:2259-2320 9347:2321-2341 9286:2342-2370 9371:2371-3110 9345:3111-3156 9365:3157-3164 9357:3165-3188 9344:3189-3274 9320:3275-3298 9356:3299-3351 9253:3352-3375 9370:3376-3518 9364:3519-3554 9355:3555-3570 9342:3571-3594 9340:3595-3744] [-1, 9265, 9266, 9377, 9374, 9369, 9352, 9242, 9243, 9244, 9245, 9373, 9367, 9376, 9349, 9372, 9360, 9347, 9286, 9371, 9345, 9365, 9357, 9344, 9320, 9356, 9253, 9370, 9364, 9355, 9342, 9340, -2]

TCAGTATCGAAATATCGGAAAAACAACGACTACGACAGGACTATGTCAACGAAATGCTCGTTGAACTACGTTGTTGAGAAGACAAGTTATTAGTTTTCTACGTTATGTGTATTATTATACGTGTTTTTTGTTTGTGCATCATACAAAGTAAATTGTGAATTTTCTTAATATATAGCTCTTCACTTCAAGCCGACACTTTTTTGTATTCGAGCTTAACAATGGGAAGAAAGAAAGGTAATAAACCACAACCAAATCCTGATGGTTCGCAACCAGCGCCATCACAGAGTAGCCAGCAGGATAGCGATACTCCCACTCGGACATCGCAAGGAACTTCTGTCTCTGAATCTCATCAAGGTACAGGACCACCTCGGGGATCGTTTTCCGGCACACCACAGCATCATTATCGAGGAAGCCAAGGACATGGTTCTGGTCAGCGTTCTGATTTCCGGGAAGCTCGTTCCTGGGGTACTGGAGCACGTTCAGATGGTTCTCACGGTCAACGTTCAGATTATAGAAGATTTCCCGCGAACAGAGCGCAGCCTGGTGGTAATCAGTATGAAAGAAGGCAACGTTCTGATTTCCAGGAACCTCGTTCCCGGGGTTATGGGCCGCGTCCAGATGGTTCTCGTGGCCAACATGCGGATTATCGAAGACCTGCCGCGGATAGAGGAGAACCTAGTGATAATGCTTCTCAAAGCCCCAGAGAATCGAGTATTAGTACTCCTCAGAAACCAGCAACTTCATCGATTTCGGCATGGGGTTCACCATCGAAACCAGCTTTTAGTCGCATGGAACATTCTGCATCTGTTAGAGATGAACCTCAGGCTAAACCAACAACTTCGAGTTGGGGAGTAAAGTATAATATTCCAGCAGGAAAGGAGCGCTCTGTTTCATCTGGTGTCTCAGAAGAGCGTTCTAAACGGACTCCTGTTGAAGAGTCTGCTTTCTCCAAACTGCCAGAAGATGCTCCAAAAGCTGGTCATAAGAGAGGTAAGAGTGAGAAAGGCCCGAGTAAGAAAAGCCCTGCTGGCAATGAAACGAACGCATCTTCCTCTAAACAACCCGTCAGTGAAAAACCGAAGGCATCATCTTCAAAGGAACAACCTACTACTGAAGATACCACAAAAATGATGGAAAACGTAGATTTGAATGAATTCCCTATGCCTGTTAGAAAGAAAAAAGAAATTAAACACACGAGTGATTACATGCTAAAGAAGATAAGCCTTGAAGTTAATTACTTGACTATTGACATCAAGACGTCTAATTTGCAAATATATCATTATGATGTGAACTTTTCGCCGGAAGCGGCTAAATATCTTTTCAGACCAGCTATGGTAATATTTGTGAATCGACATGCTCCCGGACGGCATCCGGCATTTGATGGTCGAAGAAATCTCTACGCTGCTAGTAATCCATTACCATTTAATAATATCACTGATGAGGTTACGATATTGGATGAAGTAAACCTAAGAGAAAGGAGGTATGAAATAACAGTTCAGCTCGTAGATCAACAACTGAACTTAGATGGATTAAGAGAGTATCTACGTAATGGTAGTTCTATGGATCAGCCTCAAGATTCATTGCAAGCGATGGATATAGCCATTCGGCAATCACTTTATTGTAGAGGATTTGTTCCATGCCGTCGTTCCTTCTTTTCACCACCAGCTGAACAAATTGATCTAACAGGAGGTCTTGAATTATGGTCTGGTCATTTTCAAAGCGTGGTGCTGGGTTGGAAACCATATCTAAATGTAGATGTTGCCCACAAAGCTTTTTATAAGAAACAAACTATATTAGAATTTCTCAACGCATCTATTGGAAATTTTGATCCGATGCAACGAATGGTAGATTGGCAAAGAGATGAATTGAAGAAACTTATTACCTACAAAATCGAATACCAAATGCCACCTGATGCTGGTATTAGAAAGTATAGAGTAGCTGGTATCGGTCAAACAGCTAGGGATACATTTCAGTTAGAAGAAAATGGAGTTACGAGGCAAATAACTGTAGAGGAGTATTTCCTCCAATACAAAAGATACAAAATACGGTATCCCAATTTACCACCTATGCGAACTCCGCGTGGTGACTTACTACCTTTGGAATTGTGCAGTATTTTGCCTAATCAACCATTCGTGCGAAAATTGAATCCTAAGCAAACGTCAGAAATGGTGCAAAATGCCGCTCAACCACCGAACGAGCGCAAGCAACGAATAGTACATGCGATTCGTAATGCCAGATTCAACGAAGACCCTTGTATTCAAGAATTTGGCGTGACGATTGGCGATCACTTCACCCAAGTTGACGCATGGTTGATGAAACCTCCGAAGATCGAGTACGCAAATAGGAATATAGCGGAAGTGAATAATGGTGTTTGGAGAACTAATGGTAAGTTCTTCCAGAACACCACATTAAGTAACTGGGCTATCATCAATACCGAGCTGAGGCAGATTAATGACAGTCATAAATGGGGTATCGTTAGAGCATTTCGAATGGCGGGAGCGGCGCTGGGATTAAATGTCGAAGAACCGATTTACGTTGGTGACTGCGGATTGAGAAGAGATGAGGATATGGTAAAGGAATTCAAAGCATTTAAAACTATGCCTAAACTTGTCGTCGTCATTATGAATAATTCGGAAGATTGTTACGGTAAAATAAAAAGAGCAGCGGAATTGAACTCAGGTGTGTTGACGCAGTGTTTGAAATTAAACACTGTGAAAAGAATGAATCCTATGACCGCTACAAATATTTTATTCAAAATAAATGCCAAATTGAATGGCACTAATCATATTTTATTGGATGCTATATGTCCGAATATATTGAAGAAACATAAAGTCATGATCGTTGGAGCTGATGTCACTCATCCATCACCGGGTGTTGAAAGCGTTTCAATTGCAGCGGTCGCCGCTAGTCACGATATACACGGATTCAAGTACAACATGACTTATCGGCTGCAAACATCTCGTATGGAAATGATTTACGATTTGAAAGAAATGATGAAGGAACAATTGAAACTTTATGAACAAAGCGTTGGTGAACTTCCAGAGAAAATACTGTACTTTCGAGACGGCGTCAGTGAAGGACAGTTTGCCCAAGTAAAACACGTCGAACTGACGGCCATCCGGAAAGCGTGTACAGAATATGGTGGCGATAAGTATATGCCGGGTACTACATTTTTGGTGGTACAGAAACGCCACCATACTCGATTCTTCCCGCCCTCGAGTGCTCAATGTTCCGATAAAAATAATAATGTATTCGCTGGTACGGTCGTCGATCAACAGATCACGCATCCGACTGAGGTGGATTTCTATATGGTTAGCCATCAGAGTATTAAAGGTACGGCCAGGCCGACTAAATATCATCTCTTATGGGATGACAATGATTTCGATTTAGAACCTTTGGAGCAGCTTACGTATTATCTGTGCCACATGTTCGCCAGATGTACGCGTTCTGTGTCCTATCCAGCACCGACATATTACGCTCATTTGGCGGCATTTAGAGGTCGCGCTTGGTGGGAAAAATTTCGTTTCGATCGGGCTTTGTTGCACGACGCTGCTGCATTGAGGAGATTCCAAGAGCAAAATGGCAATGTTGTGCTGAGAAATGCGCCCATGTTCTTCGTATGATCCTTTTCGGTGTTCTTGATCGAGTTATGATTGCTGATTAAAAATATAAAATATTTTAATTACATATAGCATTATATTCTTATTCAAATAATATATTATCTATATGATTGATTAAAAAAAAA

Mgfp_ amplified from pCAMBIA1303

AGTAAAGGAGAAGAACTTTTCACTGGAGTTGTCCCAATTCTTGTTGAATTAGATGGTGATGTTAATGGGCACAAAT

TTTCTGTCAGTGGAGAGGGTGAAGGTGATGCAACATACGGAAAACTTACCCTTAAATTTATTTGCACTACTGGAAAACTACCTGTTCCGTGGCCAACACTTGTCACTACTTTCTCTTATGGTGTTCAATGCTTTTCAAGATACCCAGATCATATGAAGCGGCACGACTTCTTCAAGAGCGCCATGCCTGAGGGATACGTGCAGGAGAGGACCATCTTCTTCAAGGACGACGGGAACTACAAGACACGTGCTGAAGTCAAGTTTGAGGGAGACACCCTCGTCAACAGGATCGAGCTTAAGGGAATCGATTTCAAGGAGGACGGAAACATCCTCGGCCACAAGTTGGAATACAACTACAACTCCCACAACGTATACATCATGGCCGACAAGCAAAAGAACGGCATCAAAGCCAACTTCAAGACCCGCCACAACATCGAAGACGGCGGCGTGCAACTCGCTGATCATTATCAACAAAATACTCCAATTGGCGATGGCCCTGTCCTTTTACCAGACAACCATTACCTGTCCACACAATCTGCCCTTTCGAAAGATCCCAACGAAAAGAGAGACCACATGGTCCTTCTTGAGTTTGTAACAGCTGCTGGGATTACACATGGCATGGATGAACTATACAAAG
